# GSA: An Independent Development Algorithm for Calling Copy Number and Detecting Homologous Recombination Deficiency (HRD) from Target Capture Sequencing

**DOI:** 10.1101/2021.08.02.454110

**Authors:** Dongju Chen, Minghui Shao, Pei Meng, Chunli Wang, Qi Li, Yuhang Cai, Chengcheng Song, Xi Wang, Taiping Shi

**Author notes:** Dongju Chen and Minghui Shao contributed equally to this work. Corresponding Author: Taiping Shi, BGI Genomics, BGI-Shenzhen, Shenzhen 518083, China.

## Abstract

**Background:** The gain or loss of large chromosomal regions or even whole chromosomes is termed as genomic scarring and can be observed as copy number variations resulting from the failure of DNA damage repair.

**Results:** In this study, a new algorithm called Genomic Scar Analysis (GSA) has developed and validated to calculate homologous recombination deficiency (HRD) score. The two critical submodules were tree recursion (TR) segmentation and filtering, and the estimation and correction of the tumor purity and ploidy. Then, this study evaluated the rationality of segmentation and genotype identification by the GSA algorithm and compared with other two algorithms, PureCN and ASCAT, found that the segmentation result of GSA algorithm was more logical. In addition, the results indicated that the GSA algorithm had an excellent predictive effect on tumor purity and ploidy, if the tumor purity was more than 20%. Furtherly, this study evaluated the HRD scores and *BRCA1/2* deficiency status of 195 clinical samples, and the results indicated that the accuracy was 0.98 (comparing with Affymetrix OncoScan™ assay) and the sensitivity was 95.2% (comparing with *BRCA1/2* deficiency status), both were well-behaved. Finally, HRD scores and 16 genes mutations (*TP53* and 15 HRR pathway genes) were analyzed in 17 cell lines, the results showed that there was higher frequency in HRR pathway genes in high HRD score samples.

**Conclusions:** This new algorithm, named as GSA, could effectively and accurately calculate the purity and ploidy of tumor samples through NGS data, and then reflect the degree of genomic instability and large-scale copy number variations of tumor samples.

## Background

Homologous recombination deficiency (HRD) is a functional defect in homologous recombination repair (HRR) pathway, which is responsible for repairing DNA double-stranded breaks (DSBs), and HRD is a potent tumorigenic type of DNA lesion. Tumors with HRD status are arising from germline and/or somatic mutations in *BRCA1/2* or other HRR pathway genes, promoter hypermethylation of *BRCA1* and/or *RAD51C*, and other mechanisms[1–3]. In particular, it has proved that *BRCA1* promoter hypermethylation is also a common epigenetic event in breast and ovarian cancer, ranged from 11% to 57% in different studies [4–6]. Over the last years, several studies about HRD in pan-cancer have shown that HRD occurred in many cancers with various frequency, and it was most prevalent in ovarian, breast, prostate and pancreatic cancer[7–9].

Some drugs based on DSB or single-strand break (SSB) repair have been developed and used successfully in clinical trials. Platinum salts are currently one of the most important chemotherapeutic drugs and have a broad anticancer spectrum, which introduces DNA DSBs and interstrand crosslinks. Therefore, HRD cells are supposed to be sensitive to platinum salts. The PARP family of proteins, especially PARP1, are essential for SSBs repair by the BER pathway. PARP inhibitors (PARPi) are based on inhibiting the SSBs repairing and then the accumulation of SSBs would lead to the development of fatal DSBs. Thus, PARPi are more sensitive to HRD-positive tumors through synthetic lethal interaction[10, 11]. Several studies have shown that HRD is a potential biomarker for platinum salts and PARPi in many cancers, especially in ovarian and breast cancer, and PARPi have attracted widespread attention in the targeted therapies of multiple cancers due to better efficacy and fewer side effects[12–14]. In the past years, FDA has approved niraparib and olaparib combined with bevacizumab, for specific patients with HRD-positive status, according to two critical clinical trials, QUADRA and PAOLA-1[15, 16].

Currently, HRD testing is mainly carried out by two methods. Firstly, *BRCA1/2* or other HRR pathway gene mutation detection and analysis by a custom-designed panel, but the panel gene list is various and variants of unknown significance and secondary mutations lack a standard database[17]. Secondly, identifying the results of HRD by detecting genomic damage patterns, known as genomic scars, including three biomarkers, loss of heterozygosity (LOH), telomeric allelic imbalance (TAI), and large-scale state transition (LST)[15, 16, 18]. Besides, some other methods are also developing, such as *BRCA1/RAD51C* epigenetic analysis and mutational signature analysis, *et.al.*[9], but these are still difficult for clinical application. Two commercial genomic scar detection kits have been approved by FDA, myChoice HRD CDx (Myriad Genetics, Co., Ltd.) and Foundation Focus CDx (Foundation Medicine, Co., Ltd.)[2].

Genomic scar analysis (GSA) by calculating LOH, LST and TAI scores is a dominant way to evaluate the HRD status of tumors, and the theoretical foundation of this method is that tumors with HRD phenotype would lead to large-scale copy number variation (CNV)[15, 16, 18, 19]. There are four software commonly used to detecting CNV of tumor samples, including PennCNV, ASCAT, ABSOLUTE and PureCN. PennCNV is a free software tool based on a hidden Markov model (HMM), for kilobase-resolution detection of CNVs from Illumina high-density SNP genotyping data [20]. ASCAT and ABSOLUTE were introduced to estimate tumor purity directly from SNP array data[21, 22], and both can effectively solve the impact of purity and ploidy based on SNP array data. NGS data is much higher resolution data than SNP arrays, and provides the opportunity to derive highly accurate estimates of both tumor purity and ploidy[23]. PureCN, is optimized for targeted short read sequencing data, integrates well with standard somatic variant detection pipelines, and has support for matched and/or unmatched tumor samples[24]. Currently, PureCN is a mainstream method for purity correction based on circular binary segmentation (CBS) from NGS data, but it is necessary to artificially judge the best combination from multiple combinations of purity and ploidy.

This study aims to developing and validating an algorithm, named as GSA, which can effectively and accurately calculate the purity and ploidy of tumor samples through NGS data, and then reflect the degree of genomic instability and large-scale copy number variations of tumor samples.

## Results

### The Density and Uniformity of SNP sites

The density and uniformity of SNP sites are the critical factors for the precision and resolution of CNVs identification. Herein, the capture regions distribution of HRD Panel with all chromosomes was compared with Affymetrix OncoScan™ Assay, xGen Exome Research Panel (IDT), and a pan-cancer panel (BGI) (Fig.1A). Then, chr4, chr13 and chr21 were selected randomly to compare the number and spacing of capture regions in the above four panels (Fig.1B). The Affymetrix OncoScan™ Assay is a SNP-array based on molecular inversion probe technology, a proven technology for identifying CNVs, LOH, and detecting somatic mutations. Besides, this assay could realize the high-resolution (50–300 kb) copy number detection. HRD panel is designed for detecting large-scale CNVs (at Mb level) to calculate LOH, TAI, LST, and HRD score, thus the total number of SNP sites could reduce appropriately, but not uniformity. The results showed that the SNP sites of HRD panel had similar uniformity with Affymetrix OncoScan™ Assay, but had less SNP sites (93,200 vs. 217,611). Most of the work published to date detecting CNVs is based on SNP array, thus this study defined Affymetrix OncoScan™ Assay as a correlation method to evaluate the accuracy of HRD panel based on NGS approaches[25].

**Figure 1.**
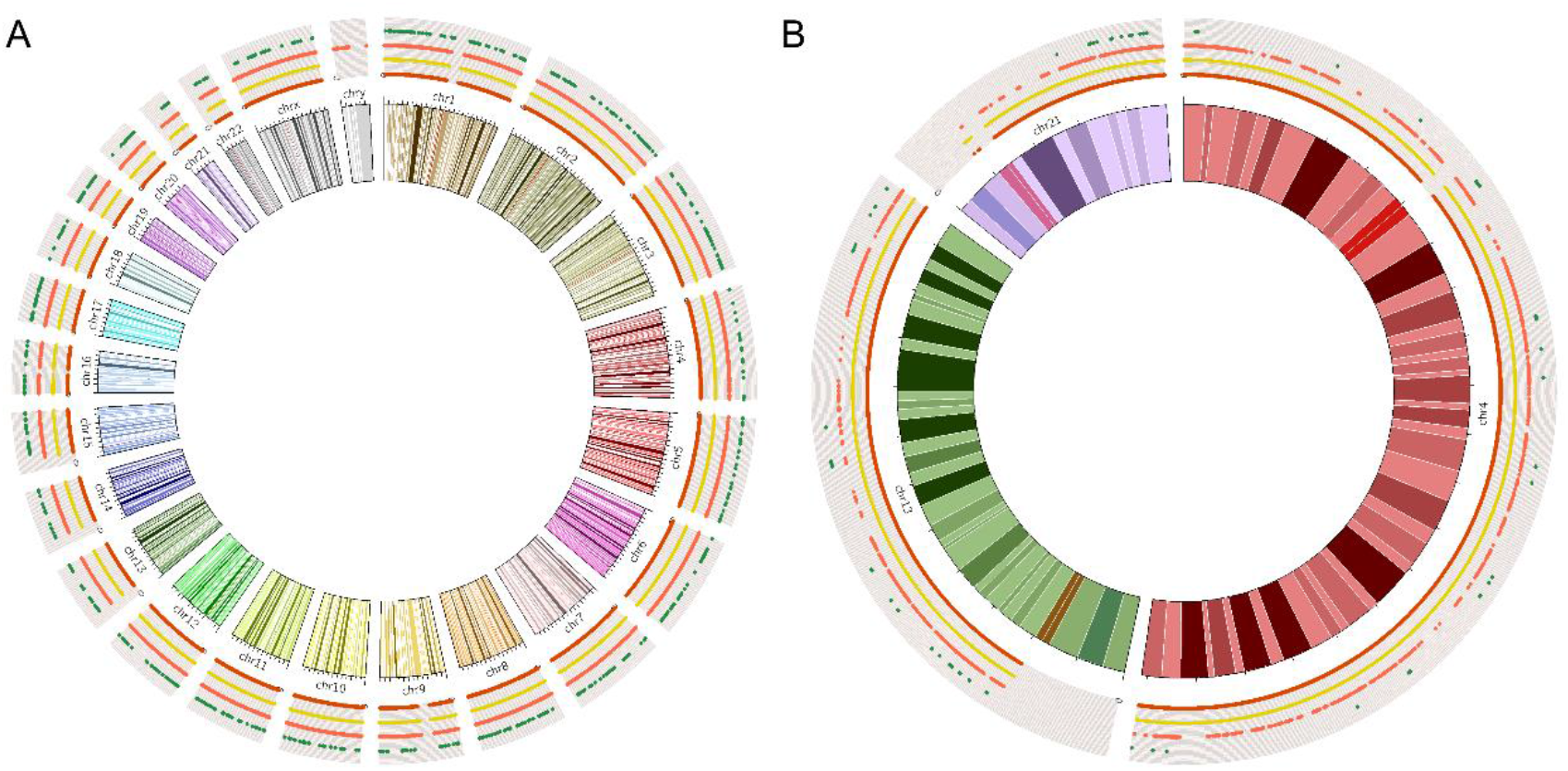
The capture regions distribution of Affymetrix OncoScan™ Assay (red line), HRD panel (yellow line), xGen Exome Research Panel (orange line), and a pan-cancer panel (green line) with all chromosomes (A) and 3 randomly selected chromosomes (B).

### The Rationality of Segmentation and Genotype Identification

Chromosome segmentation of different genotypes is the key step for detecting CNVs. This algorithm had a variety of built-in statistical methods, which could evaluate whether the adjacent chromosomal segments conform to the same genotype according to the BAF. If the adjacent sub-segmentations belonged to the same genotype, they would be combined based on circular binary segmentation (CBS). Besides, the abnormal points had been removed, thus the segmentation error rate could be reduced greatly (supplementary Fig. 4). Subsequently, the segmentation result of chromosome 2 in one patient of 195 clinical samples was selected as an example to analyze the rationality of segmentation and genotype identification (Fig.2). The results showed that the BAF and copy number of the 25M~40M region were 0.67 and 3 copies, it means that this segmentation was identified as ABB genotype. Similarly, BAF and copy number of the 115M~120M region were 1.00 and 2 copies, it means that this segmentation was identified as BB genotype. Evaluating the relationship between BAF and CNV of all segmentation were all logical.

**Figure 2.**
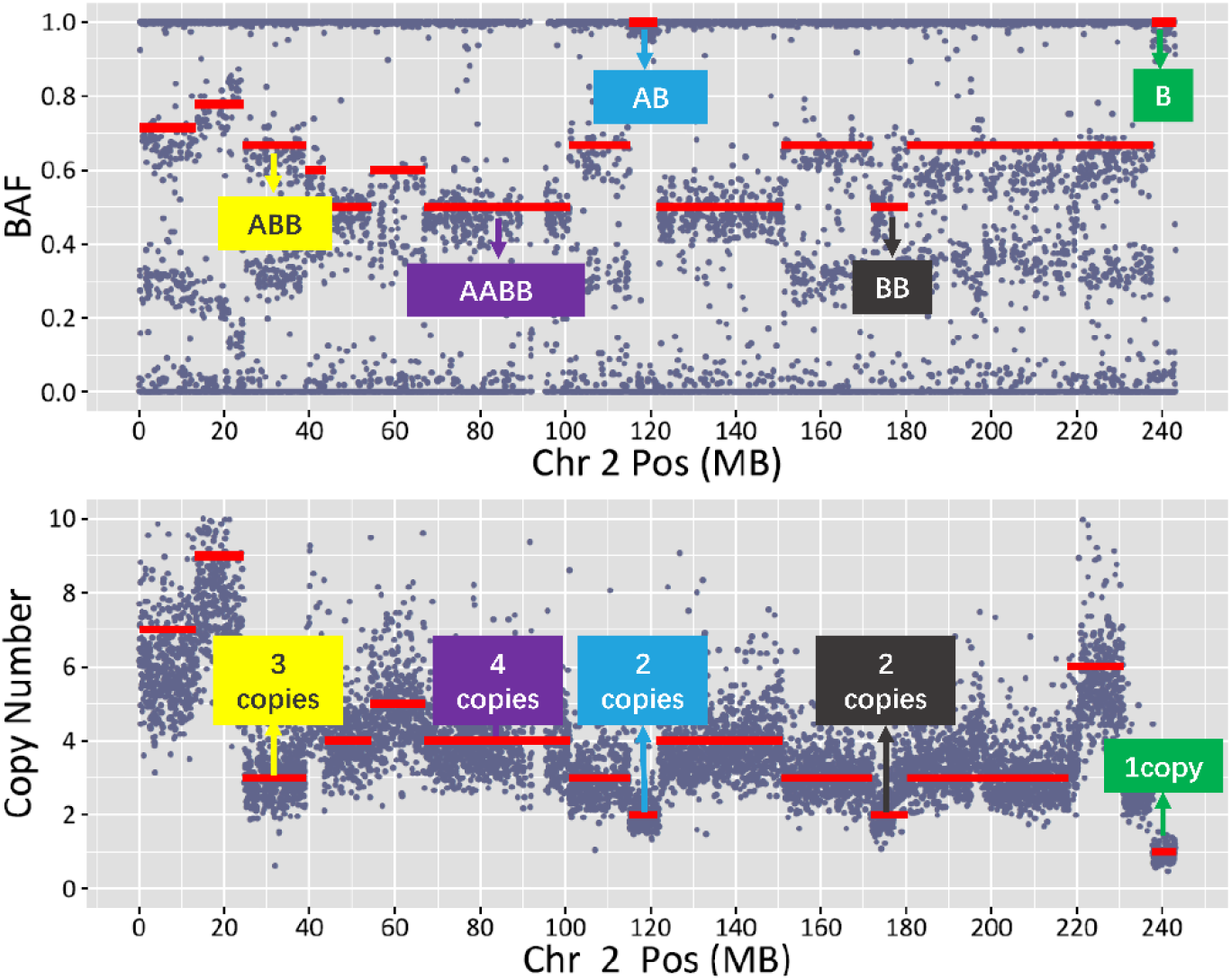
The genotype identification and the relationship between BAF and CNV in chromosome 2.

### The Self-Consistency of Purity and Ploidy Estimation

The different tumor purity samples (80%, 50%, and 20%) of HCC1143 cell line were used to evaluate the necessity and limitation of tumor purity correction. The genotype of 20M~120M in chromosome 1 of HCC1143 was considered as ABB, so the BAF distribution in this region should be characterized by a bimodal distribution with the antimode around 0.5 and peaks around 0.33 and 0.67. The density of BAF in each tumor purity was analyzed, and the results showed that the lower tumor purity, the more difficult to define ABB or AB genotype (Fig. 3A–3C). In addition, the ploidy of 195 OC and BC tumor samples had been estimated by the GSA algorithm. The ploidy ranged between 1.41 and 4.07, being characterized by a bimodal distribution with the antimode around 2.5 and peaks around 2 (near-diploid status) and 3 (near-triploid status) (Fig. 3D & Supplementary Table 3). Thus, purity and ploidy estimation were considered to the GSA algorithm. The accuracy of the GSA algorithm to predict tumor purity and ploidy were verified by the following two methods. Firstly, mix sequencing reads of different proportions of control samples into the 5 measured FFPE samples to simulate tumor purity dilution; Secondly, mix different proportions of germline DNA into 3 samples of tumor cell lines to obtain tumor cell line samples of different tumor purity. The correlation coefficient R^2^ between the theoretical tumor purity value and the actual tumor purity value calculated by GSA were 0.9813 and 0.9812, respectively, in simulated diluted tumor samples and the real diluted cell line samples (Fig. 4A, Fig. 4B, Supplementary Table 4). The tumor purity calculated by GSA maintains a good linear relationship with the theoretical purity obtained by simulated dilution, indicating that the GSA purity correction algorithm has good stability when the pathological tumor purity is higher than or equal to 20%, and the algorithm can accurately reflect the real tumor cell content. Due that tumor ploidy is an essential attribute of the sample and will not affect by the tumor purity, the smaller the fluctuation and the more accurate the ploidy correction algorithm. Similarly, the ploidy values under different tumor purity calculated by GSA are basically the same (Fig. 4C, Fig. 4D, Supplementary Table 4). In a word, GSA algorithm had an excellent predictive effect on tumor purity and tumor genome ploidy, but the pathological tumor purity should more than 20%.

**Figure 3.**
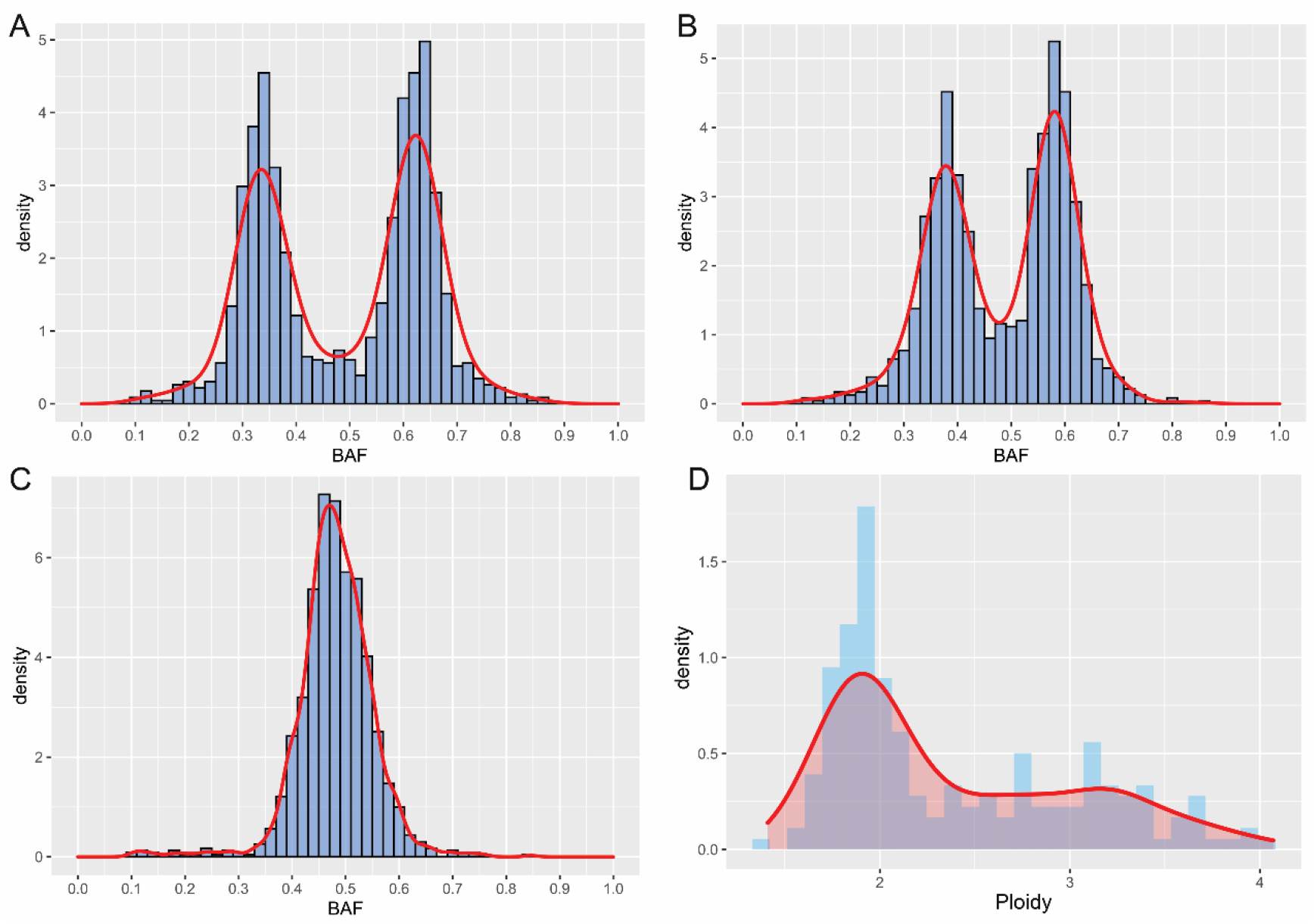
The BAF distribution of HCC1143 with 80% (A), 50% (B) and 20% (C) tumor purity in the region of 20M~120M in chromosome 1. The ploidy distribution of 195 clinical samples (D).

**Figure 4.**
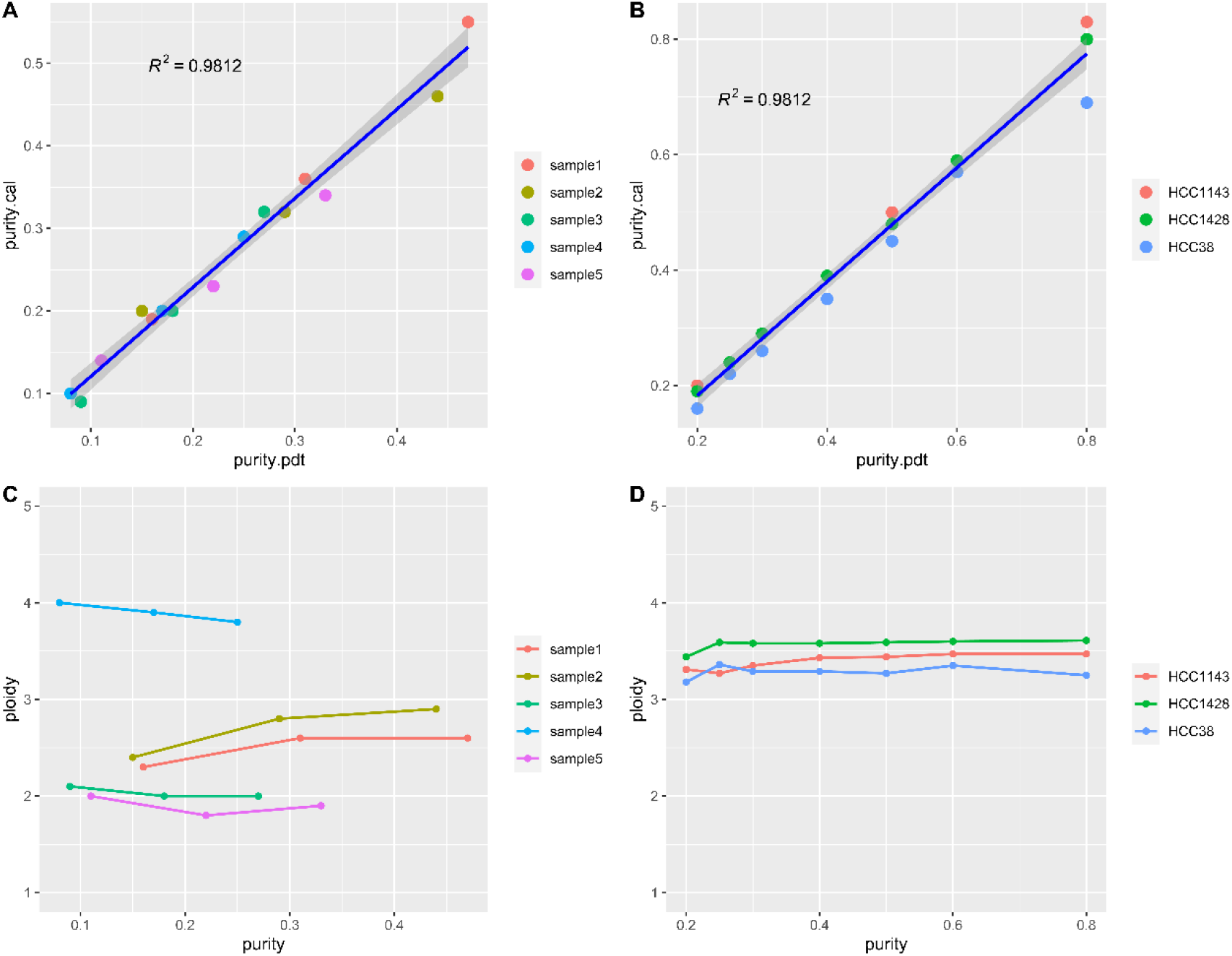
The consistency of purity and ploidy estimation with the GSA algorithm. (A) The consistency between the theoretical tumor purity value and the actual tumor purity value in the simulated diluted tumor samples. (B) The consistency between the theoretical tumor purity value and the actual tumor purity value in the real diluted cell line samples. (C) The consistency of ploidy in the simulated diluted tumor samples with different tumor purity. (D) The consistency of ploidy in the real diluted cell line samples with different tumor purity.

### The Accuracy of LOH, LST, TAI and HRD Score

Total of 40 FFPE tumor samples were both detected by Affymetrix OncoScan™ assay and the custom designed HRD Panel. BAF and LRR from NGS data and CEL file were both used to calculate HRD scores by the GSA algorithm. The results showed that the correlation coefficient of HRD scores calculated by the two method was 0.98, and the correlation coefficient of LOH, TAI and LST calculated by the two methods were 0.96, 0.89 and 0.95, respectively. It indicated that our target capture panel, containing 93,200 SNP sites, could represent the whole-genome copy number variations, and thus could generate the accurate HRD scores (Fig.5 & Table 1).

**Figure 5.**
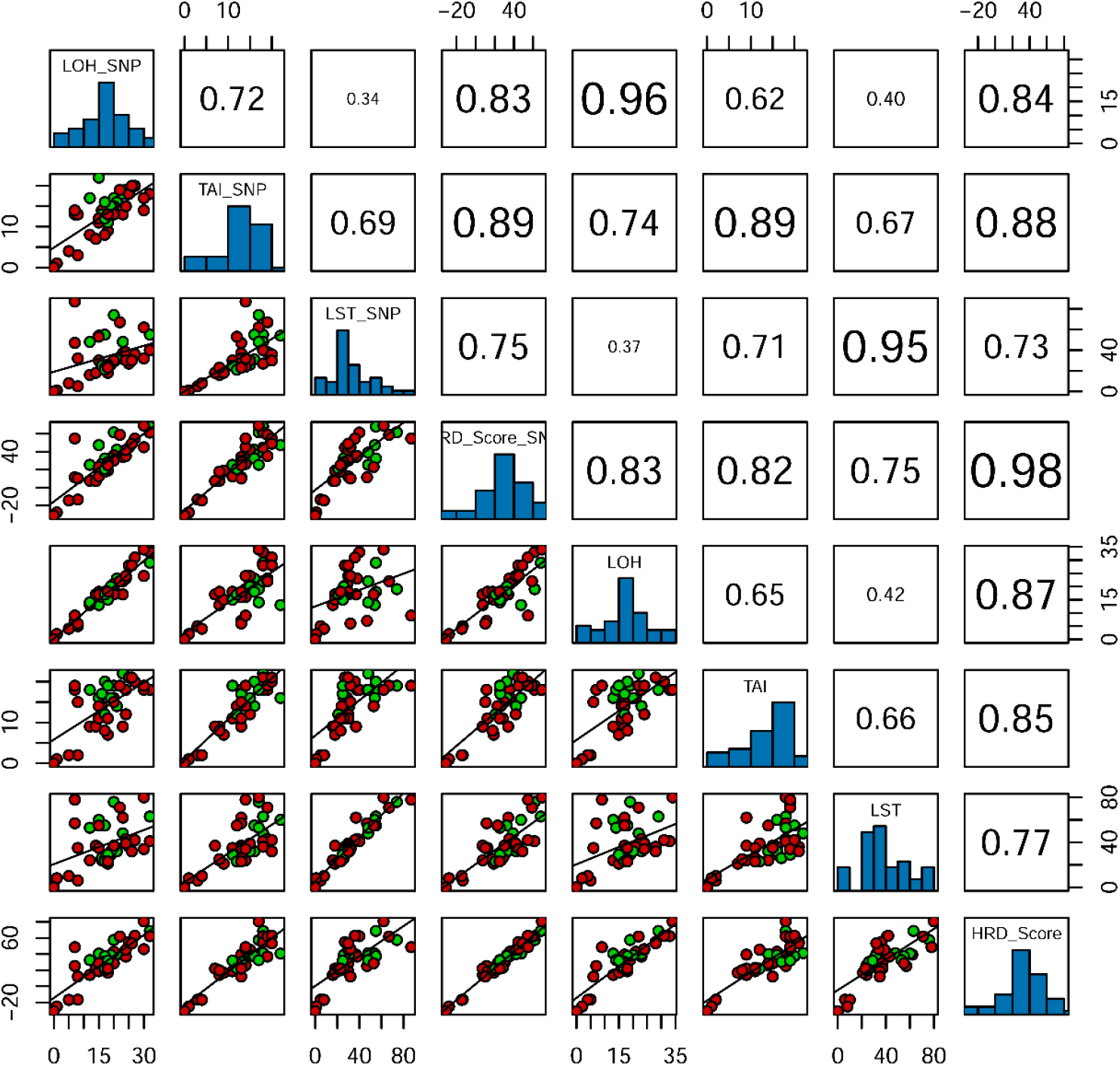
The consistency of LOH, LST, TAI and HRD score between Affymetrix OncoScan™ assay and the custom designed HRD Panel.

**Table 1.** The HRD scores of SNP array and the target panel

| Sample | SNP Array |  |  |  |  |  | HRD Panel |  |  |  |  |  |
| --- | --- | --- | --- | --- | --- | --- | --- | --- | --- | --- | --- | --- |
|  | purity | LOH | TAI | LST | ploidy | HRD Score | purity | LOH | TAI | LST | ploidy | HRD Score |
| S001 | 0.40 | 30 | 17 | 62 | 2.57 | 69.11 | 0.44 | 34 | 18 | 80 | 3.31 | 80.64 |
| S002 | 0.38 | 32 | 18 | 55 | 2.39 | 67.98 | 0.41 | 29 | 18 | 63 | 2.66 | 68.75 |
| S003 | 0.64 | 27 | 20 | 36 | 1.84 | 54.55 | 0.75 | 31 | 20 | 42 | 1.95 | 62.77 |
| S004 | 0.60 | 32 | 18 | 40 | 1.85 | 61.33 | 0.69 | 33 | 18 | 41 | 1.92 | 62.23 |
| S005 | 0.68 | 22 | 19 | 67 | 3.17 | 58.81 | 0.79 | 22 | 19 | 71 | 3.23 | 61.88 |
| S006 | 0.48 | 20 | 17 | 74 | 3.19 | 61.60 | 0.53 | 19 | 19 | 76 | 3.64 | 57.53 |
| S007 | 0.46 | 25 | 18 | 27 | 1.64 | 44.58 | 0.41 | 28 | 21 | 36 | 1.94 | 54.99 |
| S008 | 0.59 | 26 | 20 | 30 | 1.75 | 48.87 | 0.70 | 28 | 21 | 32 | 1.78 | 53.39 |
| S009 | 0.50 | 7 | 14 | 87 | 3.44 | 54.73 | 0.51 | 9 | 19 | 78 | 3.70 | 48.59 |
| S010 | 0.21 | 30 | 14 | 32 | 2.00 | 44.98 | 0.31 | 24 | 19 | 34 | 1.99 | 46.11 |
| S011 | 0.52 | 24 | 15 | 30 | 1.75 | 41.88 | 0.62 | 24 | 12 | 39 | 1.91 | 45.41 |
| S012 | 0.58 | 20 | 15 | 29 | 1.81 | 35.92 | 0.64 | 21 | 16 | 36 | 1.94 | 42.91 |
| S013 | 0.39 | 23 | 18 | 48 | 3.23 | 38.94 | 0.45 | 22 | 22 | 48 | 3.28 | 41.22 |
| S014 | 0.43 | 15 | 22 | 54 | 2.82 | 47.34 | 0.53 | 13 | 16 | 60 | 3.12 | 40.60 |
| S015 | 0.61 | 21 | 16 | 31 | 1.66 | 42.25 | 0.72 | 23 | 14 | 32 | 1.87 | 40.03 |
| S016 | 0.35 | 23 | 13 | 37 | 2.52 | 33.98 | 0.46 | 17 | 9 | 62 | 3.14 | 39.36 |
| S017 | 0.72 | 19 | 15 | 26 | 1.76 | 32.74 | 0.83 | 20 | 16 | 32 | 1.88 | 38.92 |
| S018 | 0.50 | 17 | 16 | 55 | 3.59 | 32.43 | 0.55 | 18 | 20 | 55 | 3.59 | 37.39 |
| S019 | 0.30 | 24 | 19 | 38 | 2.96 | 35.19 | 0.37 | 23 | 18 | 43 | 3.04 | 36.91 |
| S020 | 0.60 | 19 | 14 | 27 | 1.92 | 30.23 | 0.67 | 19 | 16 | 29 | 2.03 | 32.58 |
| S021 | 0.62 | 12 | 17 | 48 | 3.33 | 25.35 | 0.70 | 14 | 18 | 53 | 3.39 | 32.50 |
| S022 | 0.55 | 15 | 12 | 53 | 3.68 | 22.97 | 0.67 | 17 | 14 | 55 | 3.71 | 28.52 |
| S023 | 0.28 | 20 | 13 | 30 | 2.13 | 30.03 | 0.26 | 17 | 15 | 37 | 2.61 | 28.49 |
| S024 | 0.59 | 16 | 13 | 25 | 1.82 | 25.83 | 0.66 | 17 | 17 | 26 | 1.92 | 30.24 |
| S025 | 0.54 | 16 | 14 | 24 | 1.78 | 26.36 | 0.58 | 16 | 14 | 24 | 1.81 | 25.96 |
| S026 | 0.34 | 7 | 13 | 47 | 3.58 | 11.49 | 0.43 | 7 | 18 | 57 | 3.66 | 25.21 |
| S027 | 0.77 | 18 | 11 | 25 | 1.85 | 25.35 | 0.88 | 15 | 19 | 29 | 2.01 | 31.84 |
| S028 | 0.72 | 18 | 13 | 26 | 1.85 | 28.38 | 0.83 | 16 | 11 | 26 | 1.94 | 22.99 |
| S029 | 0.25 | 14 | 7 | 18 | 2.04 | 7.34 | 0.21 | 17 | 9 | 37 | 2.65 | 21.96 |
| S030 | 0.73 | 17 | 12 | 21 | 1.89 | 20.69 | 0.83 | 17 | 12 | 23 | 1.94 | 21.96 |
| S031 | 0.67 | 18 | 13 | 23 | 1.87 | 24.95 | 0.79 | 18 | 11 | 23 | 1.94 | 21.94 |
| S032 | 0.61 | 17 | 8 | 23 | 1.95 | 17.77 | 0.68 | 18 | 8 | 25 | 2.01 | 19.83 |
| S033 | 0.49 | 15 | 11 | 30 | 2.81 | 12.47 | 0.67 | 18 | 11 | 35 | 2.92 | 18.67 |
| S034 | 0.56 | 18 | 9 | 18 | 1.68 | 18.90 | 0.65 | 15 | 7 | 21 | 1.69 | 16.83 |
| S035 | 0.36 | 12 | 8 | 16 | 1.83 | 7.59 | 0.39 | 13 | 9 | 24 | 2.06 | 14.11 |
| S036 | 0.70 | 8 | 13 | 32 | 2.76 | 10.16 | 0.77 | 6 | 15 | 35 | 2.84 | 12.04 |
| S037 | 0.72 | 8 | 3 | 5 | 1.89 | -13.35 | 0.83 | 5 | 2 | 6 | 1.91 | -16.63 |
| S038 | 0.41 | 5 | 4 | 8 | 2.02 | -14.37 | 0.43 | 4 | 2 | 10 | 2.12 | -16.87 |
| S039 | 0.34 | 1 | 1 | 1 | 1.98 | -27.67 | 0.34 | 2 | 1 | 8 | 2.33 | -25.12 |
| S040 | 1.00 | 0 | 0 | 0 | 2.00 | -31.00 | 1.00 | 0 | 0 | 0 | 2.00 | -31.00 |

### The Relationship between HRD Score and *BRCA1/2* Deficiency Status in Clinical Samples

Deficiency of *BRCA1/2* via biallelic mutations and somatic hypermethylation (for *BRCA1*) gives rise to a deficiency status in homologous recombination repair, thus the HRD score and *BRCA1/2* deficiency status of the 195 clinical samples were all analyzed. The distribution of scores was shown for *BRCA1/2*-deficient versus *BRCA1/2*-intact samples in Figure 6. Higher HRD scores were observed in *BRCA1/2*-deficient tumors, suggesting that these tumors had a tendency towards genomic instability, and the consistency was up to 95.2% if the biological threshold was set as 30. But for other indicators, LOH, TAI and LST, there was less distinction between *BRCA1/2*-deficient tumors and *BRCA1/2*-intact tumors. Therefore, the results indicated that HRD score, combing LOH, TAI and LST, was the optimal indicator.

**Figure 6.**
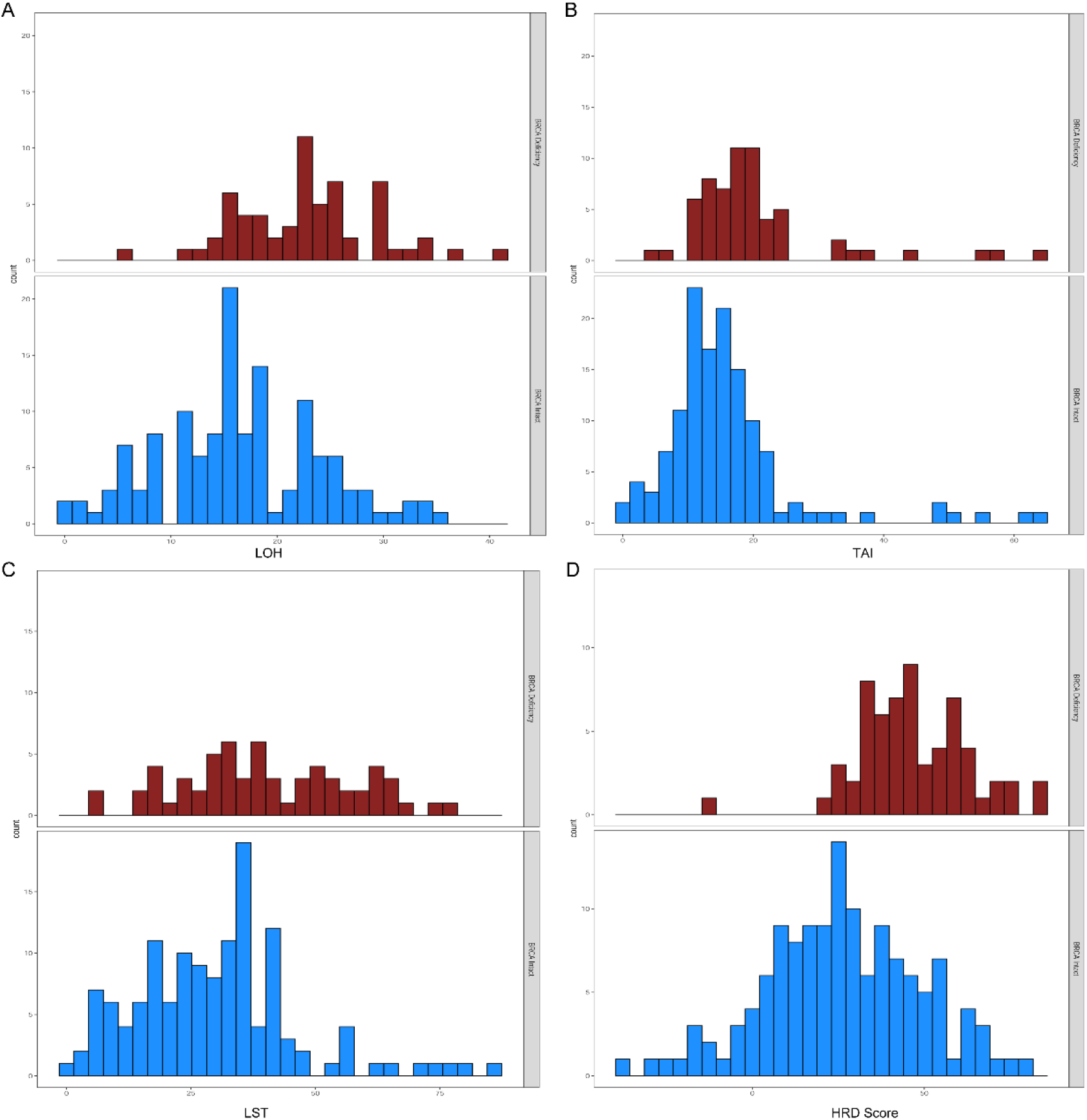
The distribution of LOH, LST, TAI and HRD score were shown as *BRCA1/2*-deficient subgroup and *BRCA1/2*-intact subgroup.

### The HRD Score and HRR Genetic Landscape of Cell lines

All the 17 ovarian and breast cancer cell lines were determined, 2 cell lines (11.76%) were m*BRCA* carriers (only including SNV and InDel variation, which are defined as likely pathogenic and pathogenic mutation carriers), and *BRCA1/2* copy number loss was detected in 8 cell lines (47.06%) (Table 2 & Fig. 7). The HRD scores distributed from −21.19 to 73.21, and the mean was 27.91, and the median was 14.42. Assuming that there is only one primary clone in the tumor samples, and amplification or loss only occur in one chromosome, the theoretic ploidy of diploid is between 1.95 and 2.04. Thus, only 1 cell line (A2780) was calculated as diploid, and other 15 cell lines (93.75%) were defined as aneuploidy (Table 2). The BAF and CN mapping of HCC38 and ZR-75-30, which had the similar ploidy correction factor but the different HRD scores, were shown as supplementary Fig.5 and supplementary Fig.6. Obviously, the genomic status of HCC38 cell line was more instable than ZR-75-30. In addition, all the cell lines with *BRCA1* methylation were showed as high HRD scores but not with any *BRCA1* mutation. Meanwhile, mutations of 13 other HRR pathway genes, *ATM*, *BARD1*, *BRIP1, CDK12, CHEK1, CHEK2, FANCL, PALB2, PPP2R2A, RAD51B, RAD51C, RAD51D, RAD54L*, and *TP53* were analyzed. *TP53*, which encodes the tumor-suppressor protein p53, was the most frequently mutated gene (82.35%), but there was no mutation in *CDK12*, *CHEK2*, *FANCL*, *PALB2*, *PPP2R2A* and *RAD51D* (Fig. 7). ZR-75-1 cell line was not detected any mutation in the above 16 genes and *BRCA1* methylation, and the HRD score was 14.42. Notably, there was higher frequency in HRR pathway genes in high HRD score samples, except for IGR-OV1, which derived from ovarian endometrioid adenocarcinoma, which is a type I epithelial ovarian cancer (EOC) and have verified as a slow growing and indolent neoplasms. The HRD score of IGR-OV1 was calculated as −14.73 but with many mutations in multiple HRR pathway genes and multi-hit mutations in *TP53* and *BRCA2*, and it indicated that this kind of ovarian cancer type might tend to genomic stability and various threshold should be set in different tumor type.

**Table 2.** The HRD scores and *BRCA1/2* mutations and methylation of 17 OC and BC cell lines

| cell line | Cancer Type | LOH | TAI | LST | ploidy | HRD Score | BRCA1/2 Mutation | BRCA1-Methylation |
| --- | --- | --- | --- | --- | --- | --- | --- | --- |
| HCC38 | BRCA | 30 | 24 | 71 | 3.34 | 73.21 | WT | 74.33% |
| HCC1806 | BRCA | 34 | 23 | 42 | 2.29 | 63.47 | WT | 1.41% |
| NCI/ADR-RES | OV | 32 | 19 | 47 | 2.32 | 61.99 | WT | 64.64% |
| HCC1143 | BRCA | 28 | 20 | 67 | 3.59 | 59.35 | WT | 1.26% |
| HCC70 | BRCA | 29 | 23 | 57 | 3.22 | 59.13 | WT | 0.28% |
| MX-1 | BRCA | 30 | 21 | 50 | 2.72 | 58.90 | BRCA1 c.2679_2682delGAAA p.K893Nfs*106 frameshift 99.84% | 2.40% |
| OVCAR4 | OV | 29 | 19 | 55 | 3.12 | 54.67 | WT | 47.56% |
| HCC1428 | BRCA | 17 | 17 | 53 | 3.62 | 30.82 | WT | 0.31% |
| ZR-75-1 | BRCA | 9 | 15 | 40 | 3.20 | 14.42 | WT | 0.17% |
| MDA-MB-453 | BRCA | 11 | 16 | 48 | 4.17 | 10.42 | WT | 0.45% |
| MDA-MB-231 | BRCA | 11 | 11 | 30 | 2.78 | 8.90 | WT | 0.19% |
| MDA-MB-361 | BRCA | 6 | 13 | 46 | 3.67 | 8.18 | WT | 1.91% |
| MDA-MB-415 | BRCA | 9 | 13 | 34 | 3.09 | 8.10 | WT | 0.09% |
| ZR-75-30 | BRCA | 7 | 13 | 41 | 3.67 | 4.10 | WT | 3.77% |
| HS-578T | BRCA | 10 | 7 | 16 | 2.47 | -5.33 | WT | 0.22% |
| IGR-OV1 | OV | 7 | 0 | 7 | 1.85 | -14.73 | BRCA1 c.1961delA p.K654Sfs*47 frameshift 50.07%<br>BRCA2 c.9448C>A p.P3150T missense 22.55%<br>BRCA2 c.9182delT p.L3061* frameshift 4.33% | 0.15% |
| A2780 | OV | 3 | 3 | 4 | 2.01 | -21.19 | WT | 0.69% |

\* BRCA represents breast cancer, and OV represents ovarian cancer.

**Figure 7.**
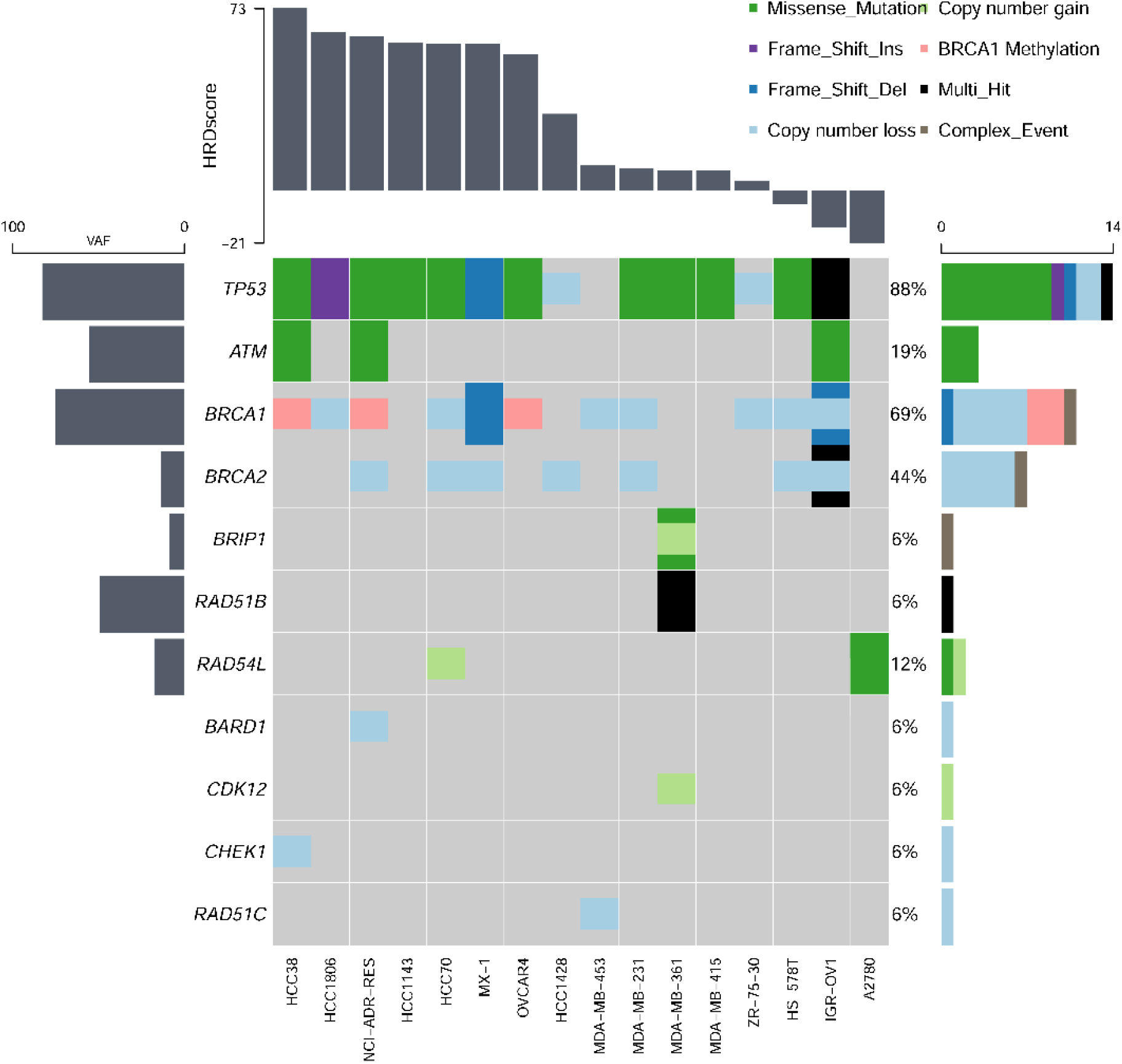
The relationship between HRD score and mutation frequency of 15 HRR pathway genes and *TP53* in 17 cell lines.

## Discussion

This study proposed a new algorithm model called GSA to detect genome aberrations based on high-throughput sequencing data, and this algorithm could accurately realize the segmentation of chromosomal regions, effectively calculate the tumor purity and tumor genome ploidy automatically and could be used for the statistical modeling of various genomic instability indicators.

Herein, the segmentation result of chromosome 2 in one clinical sample as an example to compare the chromosome segmentation effects of GSA, PureCN and ASCAT algorithms through measured samples (Supplementary Fig.7 & Supplementary Table 4). The results showed that PureCN, ASCAT and GSA algorithms divided chromosome 2 into 4, 26 and 4 fragments respectively. First of all, the BAF and copy number of the 76M~170M region were different from the adjacent segments. The BAF of this segment was concentrated around 0.5, indicating that the A/B gene ratio was closed to 1:1, and the average copy number was about 4 excluding the influence of some interference points. The result given by GSA was 4 copies, and the genotype of this fragment is AABB. But the result given by PureCN was 3 copies, which is inconsistent with BAF, and ASCAT algorithm divided the segment into 10 fragments, due to the interference of some noise points. Secondly, PureCN was disconnected at the positions of 21M, 90-92M, and 147M, but there was no significant difference at the breakpoint position judging from BAF and copy number. ASCAT divides the 10K~76M region into 7 fragments, which should belong to the same genotype inferring by the data of BAF and copy number. In addition, it should be pointed out that the GSA method has special processing for the chromosome centromere gap region. If the region distribution before and after the gap was the same, it would be merged. Overall, the GSA algorithm is more accurate for chromosome fragment segmentation.

Aneuploidy is commonly observed in cancers, and the result showed that ploidy in tumor samples is characterized by a bimodal distribution in triploid and diploid which is basically consistent with the results reported in many literatures[7]. Then, three tumor cell line samples (HCC1143, HCC1428 and HCC38) and four FFPE samples with different tumor purity were used to predict tumor purity and ploidy using GSA, ASCAT and PureCN software, respectively. The results showed that the tumor purity predicted by GSA was closest to the pathological tumor purity compared to the other two software. ASCAT has an obvious limitation in predicting samples with lower tumor purity. Meanwhile, when the actual tumor purity was low, the tumor ploidy predicted by PureCN was varies greatly. But the tumor ploidy predicted by GSA could maintain good stability (Supplementary Table 6). Although the tumor purity calculated by GSA was highly consistent with the theoretical purity of the diluted tumor cell line sample, if the tumor cell content was less than 20%, GSA algorithm could not accurately predict the tumor purity and ploidy. Normal tissue contamination and duplication of the entire genome is the most common initiation event for aneuploidy during cancer progression[3, 15, 26]. Moreover, near-tetraploid tumors show increased TAI and LST scores, but the generation of new HRD-LOH events is theoretically less likely to occur as ploidy increases[7]. The GSA algorithm added the calculation of ploidy and tumor cell purity when detecting CNVs and deducted the ploidy when calculating the HRD score. The results from clinical samples indicated that the accuracy and sensitivity of the combined HRD score calculated by GSA algorithm both were well-behaved, the accuracy was 0.98 (comparing with Affymetrix OncoScan™ assay) and the sensitivity was 95.2% (comparing with *BRCA1/2* deficiency status).

To verify whether the HRD score results obtained by the GSA algorithm can be used to assist clinical treatment efficacy, more clinical efficacy data support is needed. Unfortunately, the clinical samples used in this study lack this part of information, but we can initially compare the score of cell lines with the research results of the previous literature. The HRD scores of HCC38 and MDA-MB-231 calculated by GSA were 73.21 and 8.90, respectively. Both of the two cell lines are Claudin-low breast cancer cells, and the *BRCA* status were wildtype. In the Anne Margriet Heijink study (2019), HCC38 (GR50 = 3.6 μM) was defined as cisplatin-sensitive TNBC cell lines, and MDA-MB-231 (GR50 = 61.0 μM) was defined as cisplatin-resistant TNBC cell lines[27]. Due to the cisplatin introduces both intra- and inter-strand DNA crosslinks (ICLs), which stall replication forks and are therefore especially toxic in proliferating cells, if the homologous recombination function of cell lines is deficient, the cell lines cannot repair the double-strand breaks. Obviously, the HRD score result obtained by the GSA algorithm is consistent with the judgment of cell line drug sensitivity of the previous study[27]. With more and more application of PARP inhibitor in the clinic, we need data on the efficacy of PARP inhibitors to validate the accuracy of GSA algorithm in the real-world data.

## Conclusions

This new algorithm, named as GSA, could effectively and accurately calculate the purity and ploidy of tumor samples through NGS data, and then reflect the degree of genomic instability and large-scale copy number variations of tumor samples.

## Methods

### Tumor samples and cell lines

Archival FFPE tumor tissues were obtained from 195 ovarian and breast cancer patients who had signed the informed consents, and the study was approved by the Institutional Review Board of BGI Co., Ltd. 17 human cancer cells (HCC38, HCC1428, HCC1143, HCC1806, MX-1, HCC70, ZR-75-1, MDA-MB-453, MDA-MB-231, MDA-MB-361, MDA-MB-415, ZR-75-30, HS-578T, IGR-OV1, A2780, NCI/ADR-RES and OVCAR-4) and 3 matched-wildtype cell lines (HCC38BL, HCC1143BL and HCC1428BL) were purchased from the CoBioer Biosciences Co., Ltd. All cell lines have been verified by STR and were provided in the form of DNA status.

### Custom design HRD Panel

Variant detection requires comprehensive consideration of the detection frequency and coverage of SNP sites, and relies on statistical analysis based on the relationship between adjacent points to determine the position of fragment breakpoints and eliminate test deviations. Therefore, the principles of probe design mainly include: (1) The target region should contain high-frequency SNP sites of the population; (2) Ensure the capture region has a certain density in the whole genome; (3) Ensure that the regions are as even as possible; (4)Ensure that the target probe is synthesizable (some region probes cannot be synthesized by the supplier due to complex structure or multiple alignments lead to poor specificity); (5)Ensure that the target regions have good capture efficiency and coverage, and no obvious regional preference. Based on the above principles, a total of 93,200 high-frequency SNP loci (frequency >=5%) from the 1000 Genome Database were screened out, which are evenly distributed on each chromosome (except Y chromosomes and mitochondria).

### Library Preparation, Hybridization capture and sequencing

DNA from FFPE tissues was extracted by QIAAMP DNA FFPE TISSUE KIT (Qiagen, Hilden, Germany) according to the manufacturer’s standard protocol. Briefly, 400ng genomic DNA is fragmented and end-repaired, and a linker with a tag sequence is added to both ends of the DNA by ligase, followed by PCR amplification to form a pre-PCR library. The target DNA fragment in the library is hybridized with a combined probe containing 93,200 SNP sites and additional 2,228 capture beds targeting the complete coding region of *ATM*, *BRCA1*, *BRCA2*, *BRIP1*, *BARD1*, *CDK12*, *CHEK1*, *CHEK2*, *FANCL*, *PALB2*, *PPP2R2A*, *RAD51B*, *RAD51C*, *RAD51D*, *RAD54L* and *TP53*, *et.al*. After purification, the enriched DNA is specifically captured and amplified by PCR to obtain a post-PCR library. The post-PCR library undergoes single-strand separation, circularization and rolling circle replication to generate DNA nano balls (DNB) and sequencing was performed with 2×101 bp paired-end reads on MGISEQ-2000 platform (MGI Tech Co., Ltd.).

### Raw data quality control

Sequencing data needs to pass the basic standards of quality checks. Raw data quality control includes quality metrics for per-base sequence quality, sequence content, GC content and sequence length distribution, relative percentages of unmatched indices. Usually, the quality control parameters are set as Q30>=90%, and 40%-50% GC content.

### Data pre-processing

Raw paired-end reads were subjected to SOAPnuke (v2.0) processing to remove sequencing adapters and low-quality reads. High-quality reads were aligned to the reference human genome (GRCh37.p13), using the BWA sequence alignment software (0.7.17-r1188). PCR deduplication was performed using Picard. Average sequencing depths for tumors samples were >=150×. For each sample, SNVs were called from BAM files using an in-house software, termed as Somatk. B allele frequency (BAF) and Log R ratio (LRR) were obtained from each capture region. BAF represents the median SNP genotype frequency of each capture region, and LRR represents the normalized depth ratio of the tumor and the normal sample (or blood cell control set) in each capture region after GC-bias correction.

### TR Segmentation and Filtering Algorithm

The Tree recursion (TR) Segmentation and Filtering Algorithm was developed by C++. The input data format of the algorithm is (i) BAF data and (ii) LRR data. To reduce the noise in the input data, both BAF and LRR are preprocessed by a specially designed segmentation and filtering algorithm. First, if BAF >= 0.95 or BAF <= 0.05, defined as homozygous, the data would be removed from the BAF track because of its uselessness. Then the remaining BAF value is mirrored and flipped upward with 0.5 as the center, thus BAF= |BAF-0.5|+0.5. For LRR, the bin LRR values are also first optionally filtered for outliers, defined as the total probability density is below the 30% quantile in all bins.

Next, the in-house TR segmentation algorithm, based on the calculation of the run-length, was used to roughly segment each chromosome, as shown in supplementary Fig. 1. In this algorithm, the whole chromosome is taken as the root node, all the segmented sub-nodes are taken as the child nodes. The segmentation process can be simply described as the following steps:

a. Calculate the cumulative run-length of data (here refers to BAF and LRR) deviated from the mean 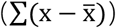, and select its maximum and minimum points as candidate breakpoints.
b. Make appropriate trade-offs of candidate breakpoints according to the location of breakpoints, length of segments, number of data points in segments, etc., that is, determine whether breakpoints should be recorded.
c. If none of the subfragment of the current fragment satisfies the record condition, a recursive judgment is initiated. Otherwise, it recursively slices its last child node.
d. After the termination condition is reached, recursion is carried out on horizontal child nodes.
e. If all child objects have been processed, the parent’s level object will continue to be processed until it is finally traced back to the root node and no new child objects are created.

Then the fragments are merged in a cyclic manner. Firstly, for each segmented fragment of chromosome was traversed by the kernel density estimation, to find out the two fragments, which are closest to the same distribution and combine them. Secondly, the statistics of the newly merged fragment and its adjacent fragments are recalculated until all indicators meet the requirements. Besides, segmentation of BAF and LRR is carried out separately, and then the union set of the merged BAF and LRR segmentation list is taken, but the regions with too short or insufficient data points are iteratively removed.

### Purity and ploidy estimation

BAF and LRR are expressed by a given genomic location as functions of the allele-specific copy numbers nA and nB, where n_A_ denotes the number of copies of the A allele and n_B_ denotes the number of copies of the B allele. Assuming tumor cell purity (p) was 1, BAF and LRR are calculated by:

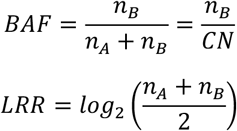

Considering the influence of nonaberrant cells in real world tumor samples and assuming that the nonaberrant cells have a total copy number of 2 for all loci, tumor ploidy correction factor (scale_factor), tumor purity (p), the measured CN (CN*) and the measured BAF (BAF*) of the FFPE samples satisfy the following relationship (Supplementary Table 1 & Supplementary Table 2).

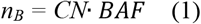

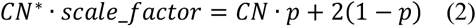

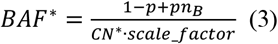

Based on Eqs (1), (2) and (3), Tumor purity can be expressed as below.

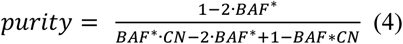

Based on the segmented chromosome fragments, the mean of all BAF value in the fragments, the percentage ranking of the BAF mean in all fragments, the theoretical CN (CN=2^LRR^*2) and the percentage ranking of the CN in all fragments are calculated. Subsequently, using the density-based scan (DBSCAN) algorithm to perform density clustering on the BAF mean-CN percentage ranking data, the chromosome fragments of the same genotype are clustered into a cluster.

For fragments of the same genotype, the calculated purity value can be approximately regarded as conforming to a normal distribution with the theoretical mean value of purity, that is Eqs (5).

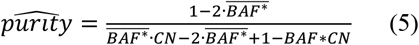

Therefore, the tumor purity is calculated by clustering the chromosome fragments of the same genotype, and bringing the measured mean value of BAF*, theoretical BAF and CN values of the specific genotype cluster into the Eqs (5).

In addition, the ploidy value of the entire genome of the sample is the weighted average of the copy number of each segment of the chromosome.

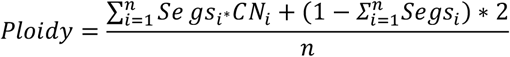

Segs_i_ is the proportion of each segment on the reference genome, and CNi is the calculated copy number of the segment.

### Calculation of LOH, TAI, and LST scores

HRD-LOH score was defined as the number of LOH regions longer than 15 Mb. HRD-TAI score was defined as the number of regions with allelic imbalance that (a) extend to one of the subtelomeres, (b) do not cross the centromere and (c) are longer than 11 Mb. HRD-LST score is the number of break points between regions longer than 10 Mb after filtering out regions shorter than 3 Mb.

Aneuploidy is a common event in cancer patients, so more copy number variations will be detected by the high-throughput sequencing data. However, these copy number abnormalities may not be caused by the failure of homologous recombination repair, it will make the final HRD score calculation biased. Calculating accurately HRD scores depend on BAF and copy number, but the aneuploidy properties and various purity of tumor samples will affect the actual value of BAF and copy number. Thus, it is necessary to make appropriate correction, and the calculation formula of HRD score is preliminarily determined as follows:

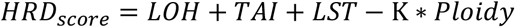

Here, K is the coefficient of correction, which is a constant. Besides, the whole analysis flowchart was shown as Supplementary Fig. 2. However, the constant depends on the type of cancer, sample type, target region size, and sequencing platform, et.al. This study screened 62 BRCA1/2-deficiency samples and 37 BRCA1/2 wildtype clinical samples of the 195 patients to explore the reasonable constant K. Finally, when the correction coefficient was set as 15.5, the AUC of the model is 88.3%, the sensitivity is 95.2%, and the threshold is 30 (Supplementary Fig. 3).

### *BRCA1/2* and other HRR gene mutation analysis

Variants were named according to HGVS (Human Genome Variation Society; http://www.hgvs.org/). Point mutations, short InDels, copy number variants were identified from NGS data, and interpreted in accordance with the “Genetic Variation Annotation Standards and Guidelines” (2015 Edition) issued by the American College of Medical Genetics (ACMG) for germline mutation, and the “Cancer mutation interpretation of guidelines and standards (2017 Edition)” for somatic mutation, respectively. *BRCA1/2* locus-specific loss of heterozygosity were analyzed as follows: a) if the mutation frequency of the SNP on the control sample is between 35% and 65%, it is recorded as a heterozygous mutation; b) if the mutation frequency of the SNP on the tumor sample is greater than 65% or less than 35%, it is recorded as a homozygous mutation; c) if the mutation frequency of a SNP site meets the both conditions, the SNP site is marked as a LOH site, otherwise it is marked as a non-LOH site; d) if the number of LOH sites on *BRCA1/2* is greater than the number of non-LOH sites, it is considered that LOH has occurred in *BRCA1/2*.

### *BRCA1* Promoter methylation quantitative PCR assays

DNA methylation-sensitive and methylation-dependent restriction enzymes were used to selectively digest unmethylated or methylated genomic DNA, respectively. Post-digest DNA was quantified by real-time PCR using a 344-bp PCR-generated primer that spanned *BRCA1* exon 1. The relative concentrations of differentially methylated DNA are determined by comparing the amount of each digest with that of a mock digest. A cutoff of 10% was used to define samples as “methylated”.

### The definition of *BRCA1/2*-deficiency

*BRCA1/2*-deficiency is defined as either (i) one deleterious mutation in *BRCA1* or *BRCA2*, with LOH in the wild-type copy or (ii) two deleterious mutations in the same gene or (iii) promoter methylation of *BRCA1* with LOH in the wild-type copy.

### Affymetrix OncoScan™ Assay

The Affymetrix OncoScan™ assay utilizes the Molecular Inversion Probe (MIP) assay technology for the detection of SNP genotyping, and has subsequently been used for identifying other types of genetic variation including focal insertions and deletions, large fragment CNV, LOH, and even somatic mutation. This assay has been shown over time to perform well with highly degraded DNA, such as that derived from FFPE- preserved tumor samples of various ages and with <100 ng DNA of starting material, thus making the assay a natural choice in cancer clinical research. This assay captured the alleles of 217,611 SNPs and then the original CEL files were obtained by Affymetrix Genechip Scanner were converted to the OSCHP files by Chromosome Analysis Suite 3.0.

### Statistical analysis

All statistical analysis was conducted using R version 3.6.1 (R Core Team, 2013) with an α of 0.05. The statistical tools employed in this study include Student’s t-test and one-way ANOVA analysis of variance. All reported *P* values were two-sided. *P* <0.05 was considered to be statistically significant. The Pearson correlation were used to evaluating the consistency of two different methods. The two-dimensional normal distribution function was used to remove outliers. Same distribution statistical test was used to compare the difference between adjacent fragments. DBSCAN density clustering algorithm was used to identify different genotypes.

## List of abbreviations

NGS: next generation sequencing
GSA: Genomic Scar Analysis
HRD: homologous recombination deficiency
HRR: homologous recombination repair
DSB: double-stranded break
SSB: single-strand break
PARPi: PARP inhibitors
LOH: loss of heterozygosity
TAI: telomeric allelic imbalance
LST: large-scale state transition
CNV: copy number variation
HMM: hidden Markov model
CBS: circular binary segmentation
BAF: B allele frequency
LRR: Log R ratio
TR: tree recursion

## Declarations

### Ethics approval and consent to participate

All patients had signed the informed consents, and the study was approved by the Institutional Review Board of BGI Co., Ltd.

### Consent for publication

Not applicable.

### Availability of data and materials

The data generated during this study are also available in the CNGB Nucleotide Sequence Archive (CNSA: https://db.cngb.org/cnsa; accession number CNP0002015).

All data analyzed during this study are included in this published article and its supplementary information files.

The basic information of GSA is available at https://github.com/shaominghui/GSA.

### Authors’ contributions

Dongju Chen and Minghui Shao wrote this manuscript with contributions from all authors. Minghui Shao, Pei Meng and Chunli Wang designed the GSA and analyzed the variant caller results. Dongju Chen, Qi Li, Yuhang Cai and Chengcheng Song performed the experiments and interpreted the results. Taiping Shi directed all aspects of the project from concept, to design, to engineering to experimentation. All authors have read and approved the final version of the manuscript.

