## Supplementary Material for "GSA: An Independent Development Algorithm for Calling Copy Number and Detecting Homologous Recombination Deficiency (HRD) from Target Capture Sequencing"

**Supplementary Material Captions**

Supplemental Fig. 1 The schematic diagram for tree recursion (TR) segmentation algorithm.

Supplemental Fig. 2 The bioinformation analysis flow chart of the GSA algorithm.

Supplemental Fig. 3 The ROC curve modelled by 62 *BRCA1/2*-deficiency samples and 37 *BRCA1/2* wildtype clinical samples of the 195 patients.

Supplemental Fig. 4 The comparison diagram between before and after removing the abnormal segments according to BAF and LRR. (A) Before removing the abnormal segments according to BAF. (B) After removing the abnormal segments according to BAF. (C) Before removing the abnormal segments according to LRR. (D) After removing the abnormal segments according to LRR.

Supplemental Fig. 5 The BAF and CN mapping of HCC38 with high HRD score.

Supplemental Fig. 6 The BAF and CN mapping of ZR-75-30 with low HRD score.

Supplemental Fig. 7 The chromosome segmentation effects comparison of GSA, PureCN and ASCAT algorithms in chromosome 2.

Supplementary Table 1 The relationship between theoretical BAF with different tumor purity at specific genotype.

Supplementary Table 2 The relationship between theoretical copy number with different tumor purity at specific genotype.

Supplementary Table 3 The HRD score and *BRCA1/2* deficiency status of 195 clinical samples

Supplemental Table 4 Comparison of the chromosome segmentation and copy number of GSA, PureCN, and ASCAT.

Supplemental Table 5 Tumor purity and ploidy values calculated by tumor cell lines and clinical samples with different tumor purity.

Supplemental Table 6 Comparison of the tumor purity and ploidy predicted by GSA, PureCN, and ASCAT.

Supplementary Figure 1 The schematic diagram for tree recursion (TR) segmentation algorithm.

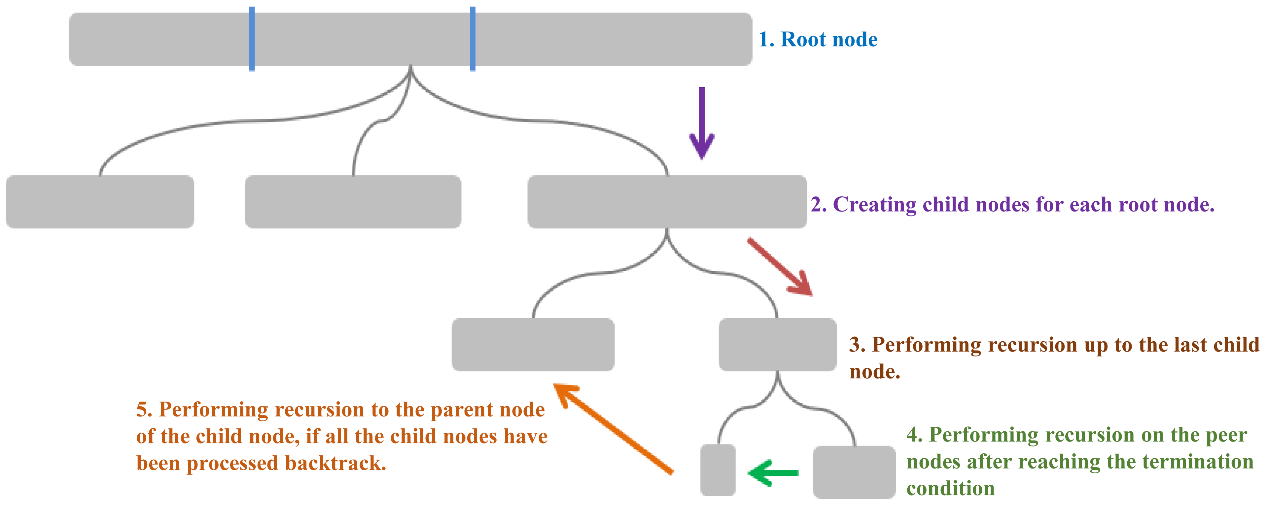

Supplemental Fig. 2 The bioinformation analysis flow chart of the GSA algorithm.

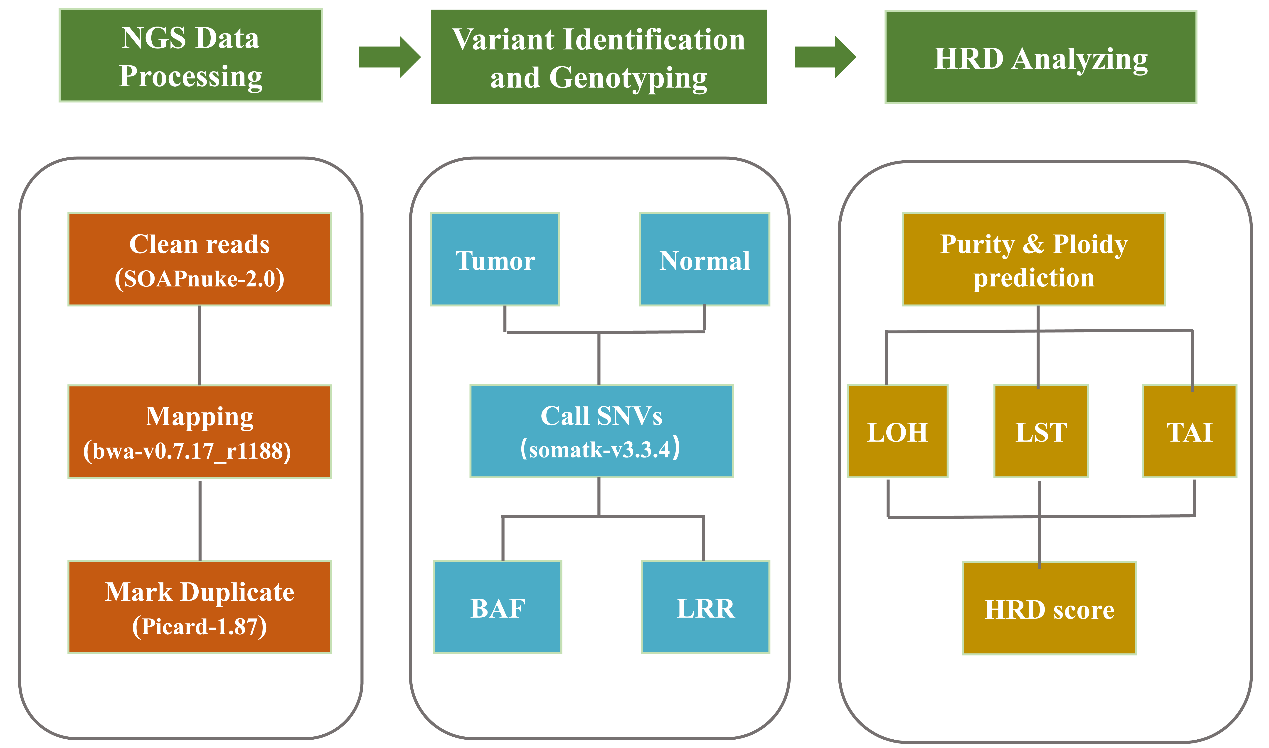

Supplemental Fig. 3 The ROC curve modelled by 62 *BRCA1/2*-deficiency samples and 37 *BRCA1/2* wildtype clinical samples of the 195 patients.

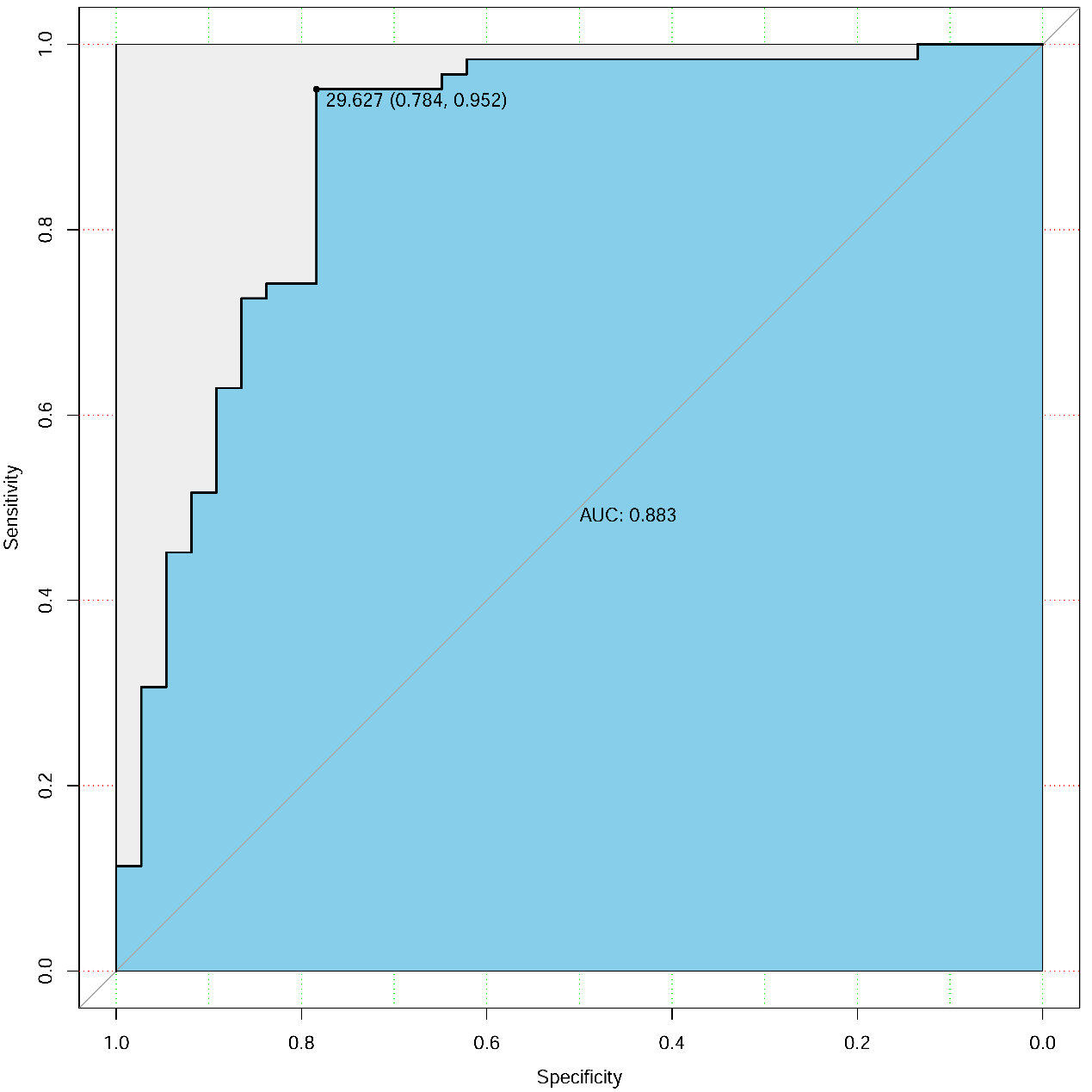

Supplemental Fig. 4 The comparison diagram between before and after removing the abnormal segments according to BAF and LRR. (A) Before removing the abnormal segments according to BAF. (B) After removing the abnormal segments according to BAF. (C) Before removing the abnormal segments according to LRR. (D) After removing the abnormal segments according to LRR.

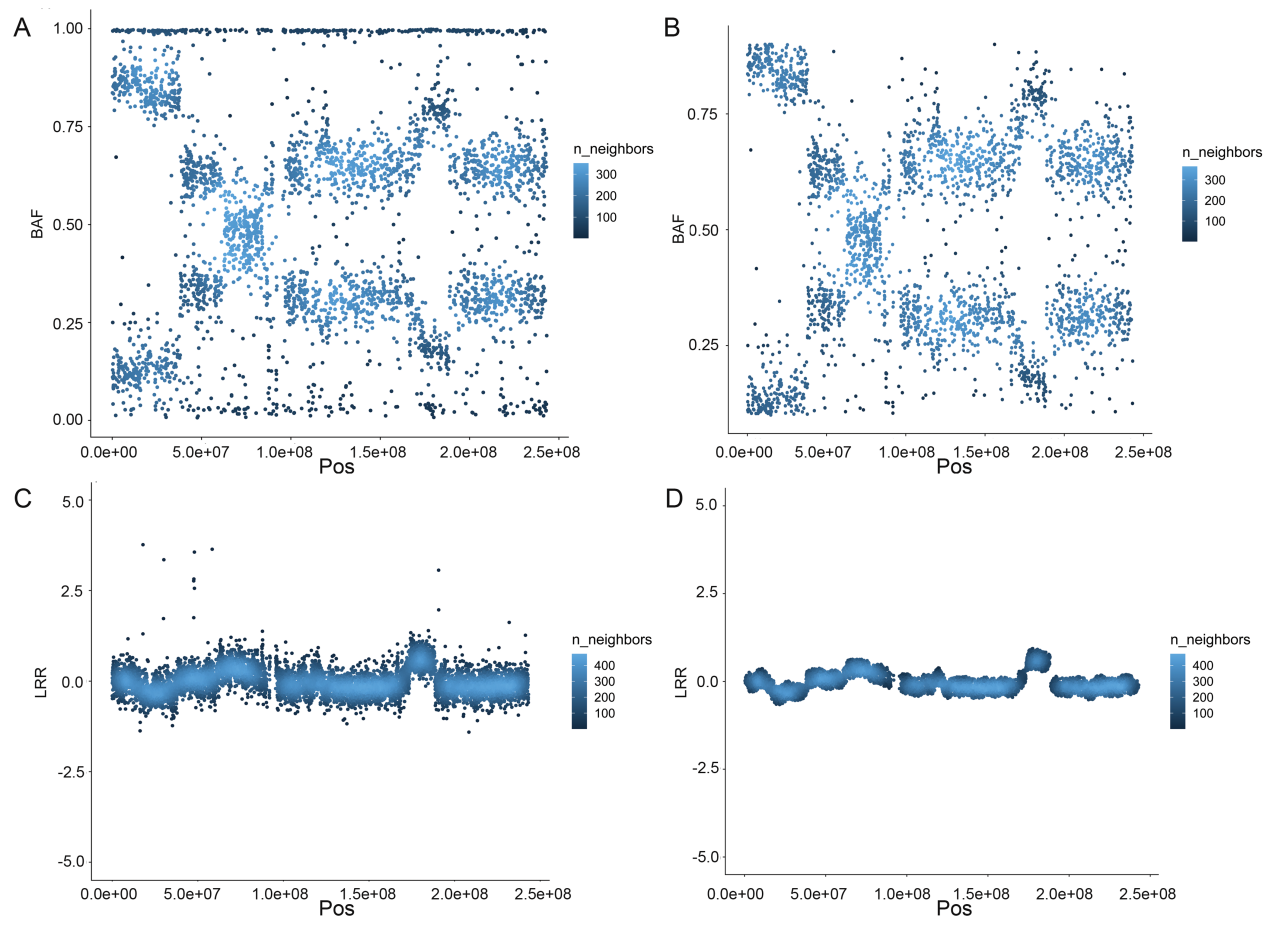

Supplemental Fig. 5 The BAF and CN mapping of HCC38 with high HRD score.

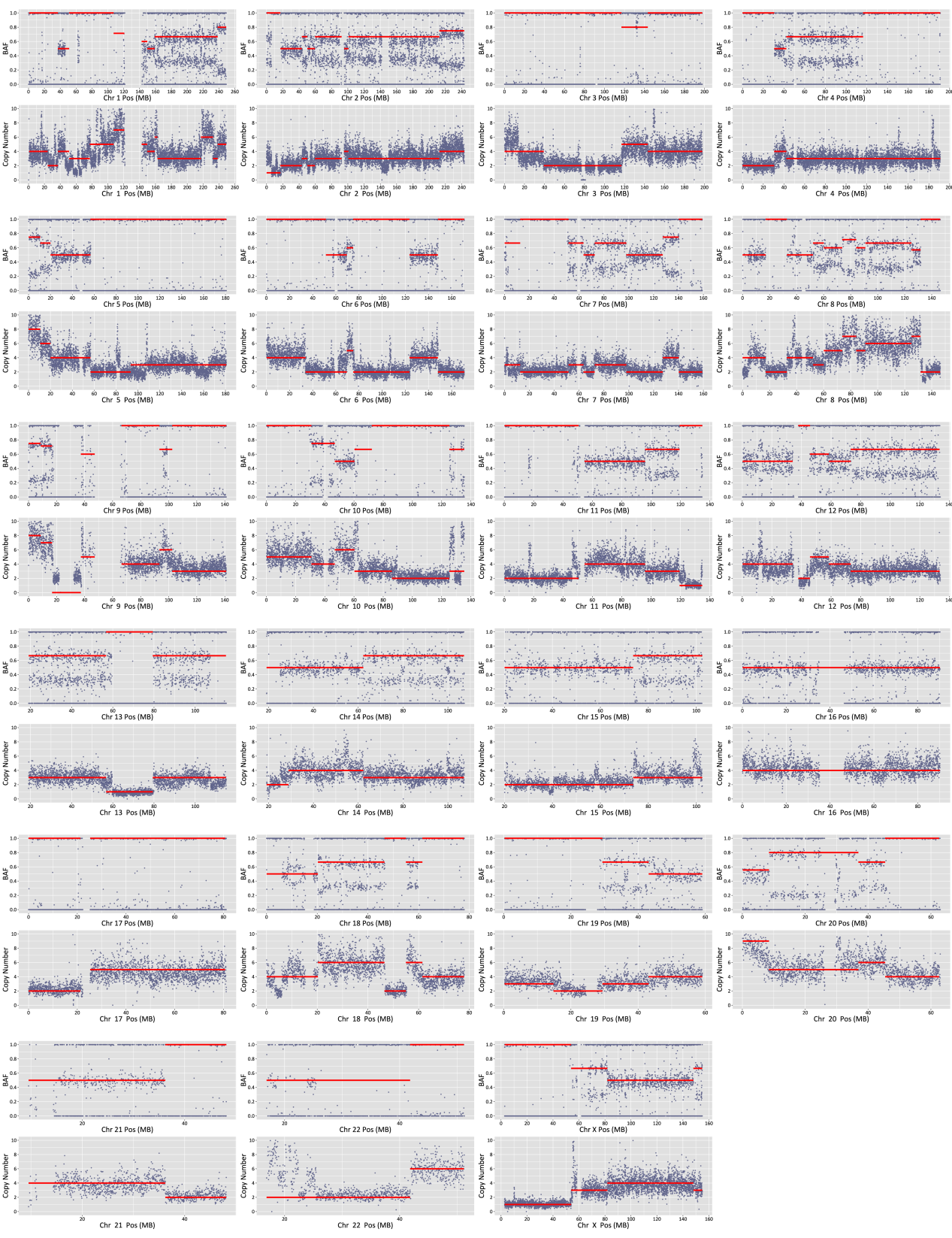

Supplemental Fig. 6 The BAF and CN mapping of ZR-75-30 with low HRD score.

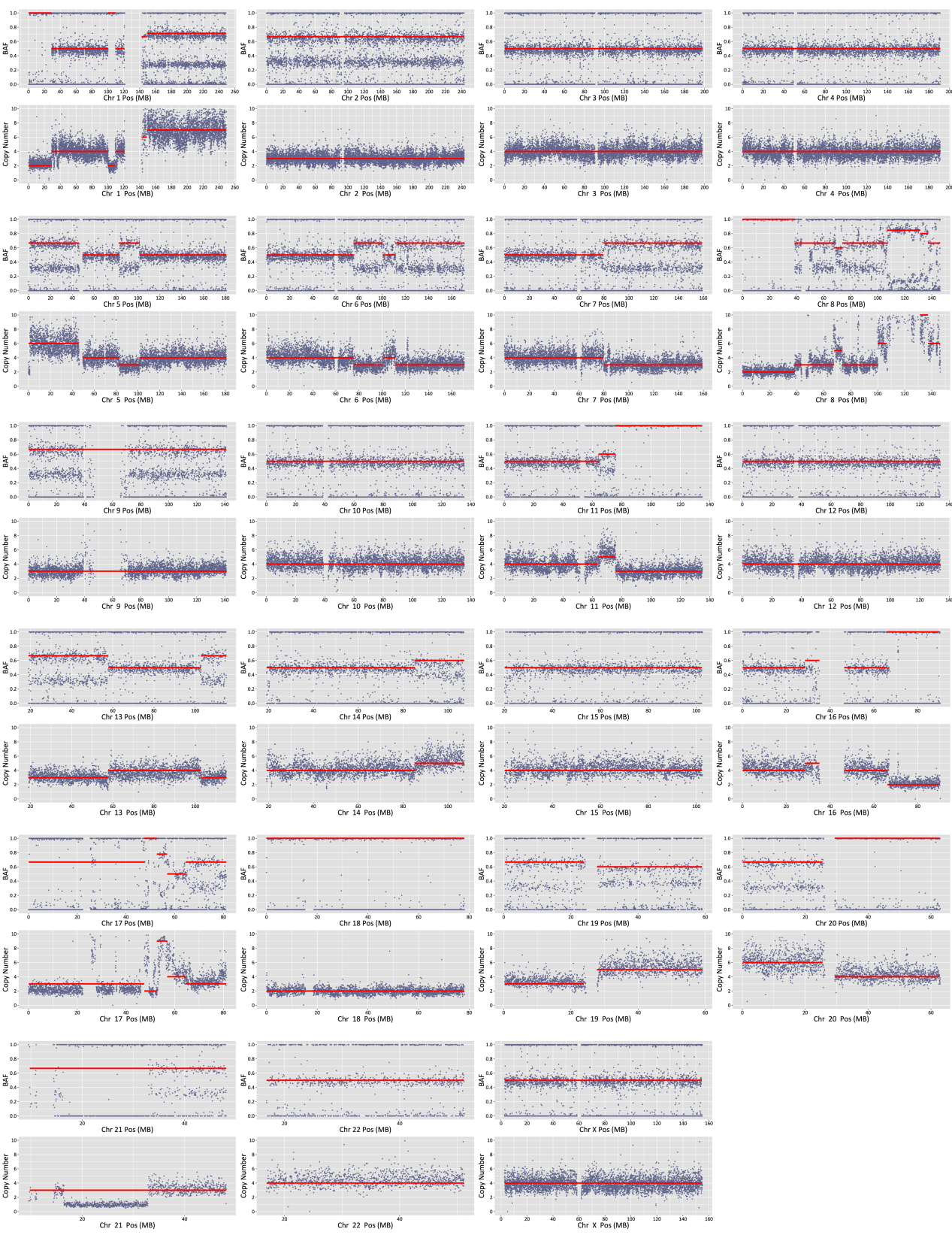

Supplemental Fig. 7 The chromosome segmentation effects comparison of GSA, PureCN and ASCAT algorithms in chromosome 2.

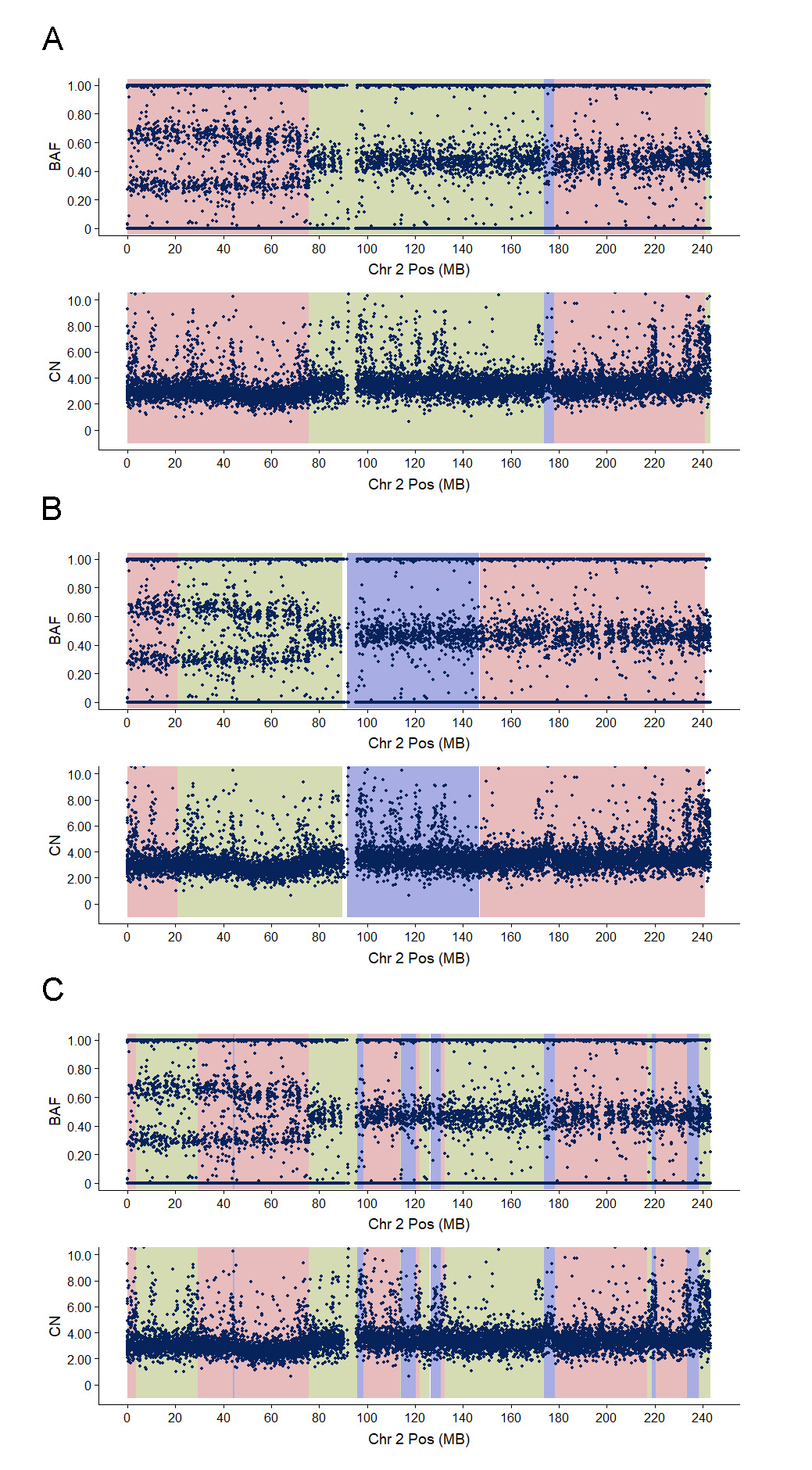

Supplementary Table 1 The relationship between theoretical BAF with different tumor purity at specific genotype.

| Tumor purity | Theoretical BAF value of a specific genotype | | | | | | | | | | |
| --- | --- | --- | --- | --- | --- | --- | --- | --- | --- | --- | --- |
|  | B | AB | BB | ABB | BBB | AABB | ABBB | BBBB | AABBB | ABBBB | BBBBB |
| 100% | 1.00 | 0.50 | 1.00 | 0.67 | 1.00 | 0.50 | 0.75 | 1.00 | 0.60 | 0.80 | 1.00 |
| 80% | 0.83 | 0.50 | 0.90 | 0.64 | 0.93 | 0.50 | 0.72 | 0.94 | 0.59 | 0.77 | 0.96 |
| 60% | 0.71 | 0.50 | 0.80 | 0.62 | 0.85 | 0.50 | 0.69 | 0.88 | 0.58 | 0.74 | 0.90 |
| 40% | 0.63 | 0.50 | 0.70 | 0.58 | 0.75 | 0.50 | 0.64 | 0.79 | 0.56 | 0.69 | 0.81 |
| 20% | 0.56 | 0.50 | 0.60 | 0.55 | 0.64 | 0.50 | 0.58 | 0.67 | 0.54 | 0.62 | 0.69 |

Supplementary Table 2 The relationship between theoretical copy number with different tumor purity at specific genotype.

| Tumor purity | Theoretical copy number of a specific genotype | | | | | | | | | | |
| --- | --- | --- | --- | --- | --- | --- | --- | --- | --- | --- | --- |
|  | B | AB | BB | ABB | BBB | AABB | ABBB | BBBB | AABBB | ABBBB | BBBBB |
| 100% | 1.0 | 2.0 | 2.0 | 2.0 | 3.0 | 4.0 | 4.0 | 4.0 | 5.0 | 5.0 | 5.0 |
| 80% | 1.2 | 2.0 | 2.0 | 2.8 | 2.8 | 3.6 | 3.6 | 3.6 | 4.4 | 4.4 | 4.4 |
| 60% | 1.4 | 2.0 | 2.0 | 2.6 | 2.6 | 3.2 | 3.2 | 3.2 | 3.8 | 3.8 | 3.8 |
| 40% | 1.6 | 2.0 | 2.0 | 2.4 | 2.4 | 2.8 | 2.8 | 2.8 | 3.2 | 3.2 | 3.2 |
| 20% | 1.8 | 2.0 | 2.0 | 2.2 | 2.2 | 2.4 | 2.4 | 2.4 | 2.6 | 2.6 | 2.6 |

Supplementary Table 3 The HRD score and *BRCA1/2* deficiency status of 195 clinical samples

| Sample | Cancer Type | Purity | LOH | TAI | LST | Ploidy | HRD Score | BRCA Deficiency Status |
| --- | --- | --- | --- | --- | --- | --- | --- | --- |
| S001 | OC | 0.44 | 34 | 18 | 80 | 3.31 | 80.70 | N |
| S002 | OC | 0.41 | 29 | 18 | 63 | 2.66 | 68.77 | Y |
| S003 | OC | 0.75 | 31 | 20 | 42 | 1.95 | 62.78 | N |
| S004 | OC | 0.69 | 33 | 18 | 41 | 1.92 | 62.24 | N |
| S005 | OC | 0.79 | 22 | 19 | 71 | 3.23 | 61.94 | N |
| S006 | OC | 0.53 | 19 | 19 | 76 | 3.64 | 57.58 | Y |
| S007 | OC | 0.41 | 28 | 21 | 36 | 1.94 | 54.93 | N |
| S008 | OC | 0.70 | 28 | 21 | 32 | 1.78 | 53.41 | N |
| S009 | OC | 0.51 | 9 | 19 | 78 | 3.70 | 48.65 | N |
| S010 | OC | 0.31 | 24 | 19 | 34 | 1.99 | 46.16 | N |
| S011 | OC | 0.62 | 24 | 12 | 39 | 1.91 | 45.40 | N |
| S012 | OC | 0.64 | 21 | 16 | 36 | 1.94 | 42.93 | N |
| S013 | OC | 0.45 | 22 | 22 | 48 | 3.28 | 41.16 | Y |
| S014 | OC | 0.53 | 13 | 16 | 60 | 3.12 | 40.64 | Y |
| S015 | OC | 0.72 | 23 | 14 | 32 | 1.87 | 40.02 | Y |
| S016 | OC | 0.46 | 17 | 9 | 62 | 3.14 | 39.33 | N |
| S017 | OC | 0.83 | 20 | 16 | 32 | 1.88 | 38.86 | Y |
| S018 | OC | 0.55 | 18 | 20 | 55 | 3.59 | 37.36 | Y |
| S019 | OC | 0.37 | 23 | 18 | 43 | 3.04 | 36.88 | N |
| S020 | OC | 0.67 | 19 | 16 | 29 | 2.03 | 32.54 | N |
| S021 | OC | 0.70 | 14 | 18 | 53 | 3.39 | 32.46 | Y |
| S022 | OC | 0.67 | 17 | 14 | 55 | 3.71 | 28.50 | N |
| S023 | OC | 0.26 | 17 | 15 | 37 | 2.61 | 28.55 | N |
| S024 | OC | 0.66 | 17 | 17 | 26 | 1.92 | 30.24 | Y |
| S025 | OC | 0.58 | 16 | 14 | 24 | 1.81 | 25.95 | N |
| S026 | OC | 0.43 | 7 | 18 | 57 | 3.66 | 25.27 | N |
| S027 | OC | 0.88 | 15 | 19 | 29 | 2.01 | 31.84 | Y |
| S028 | OC | 0.83 | 16 | 11 | 26 | 1.94 | 22.93 | N |
| S029 | OC | 0.21 | 17 | 9 | 37 | 2.65 | 21.93 | N |
| S030 | OC | 0.83 | 17 | 12 | 23 | 1.94 | 21.93 | Y |
| S031 | OC | 0.79 | 18 | 11 | 23 | 1.94 | 21.93 | N |
| S032 | OC | 0.68 | 18 | 8 | 25 | 2.01 | 19.85 | N |
| S033 | OC | 0.67 | 18 | 11 | 35 | 2.92 | 18.74 | N |
| S034 | OC | 0.65 | 15 | 7 | 21 | 1.69 | 16.81 | N |
| S035 | OC | 0.39 | 13 | 9 | 24 | 2.06 | 14.07 | N |
| S036 | OC | 0.77 | 6 | 15 | 35 | 2.84 | 11.98 | N |
| S037 | OC | 0.83 | 5 | 2 | 6 | 1.91 | -16.61 | N |
| S038 | OC | 0.43 | 4 | 2 | 10 | 2.12 | -16.86 | N |
| S039 | OC | 0.34 | 2 | 1 | 8 | 2.33 | -25.12 | N |
| S040 | OC | 1.00 | 0 | 0 | 0 | 2.00 | -31.00 | N |
| S041 | OC | 0.78 | 29 | 24 | 75 | 2.75 | 85.38 | Y |
| S042 | OC | 0.89 | 41 | 24 | 46 | 1.69 | 84.81 | Y |
| S043 | OC | 0.60 | 37 | 24 | 41 | 1.65 | 76.43 | Y |
| S044 | BC | 0.41 | 36 | 64 | 15 | 2.57 | 75.17 | N |
| S045 | OC | 0.56 | 34 | 19 | 64 | 2.71 | 75.00 | Y |
| S046 | OC | 0.63 | 34 | 49 | 17 | 1.90 | 70.55 | N |
| S047 | OC | 0.29 | 29 | 14 | 58 | 1.98 | 70.31 | Y |
| S048 | OC | 0.52 | 32 | 23 | 42 | 1.79 | 69.26 | Y |
| S049 | OC | 0.78 | 32 | 22 | 39 | 1.74 | 66.03 | N |
| S050 | OC | 0.70 | 27 | 23 | 57 | 2.66 | 65.77 | N |
| S051 | BC | 0.25 | 25 | 55 | 25 | 2.54 | 65.63 | N |
| S052 | BC | 0.60 | 25 | 64 | 24 | 3.16 | 64.02 | Y |
| S053 | OC | 0.56 | 29 | 58 | 7 | 1.94 | 63.93 | Y |
| S054 | OC | 0.79 | 34 | 21 | 35 | 1.69 | 63.81 | Y |
| S055 | OC | 0.59 | 24 | 22 | 66 | 3.16 | 63.02 | Y |
| S056 | OC | 0.65 | 26 | 11 | 86 | 3.97 | 61.47 | N |
| S057 | BC | 0.79 | 26 | 44 | 20 | 1.90 | 60.55 | Y |
| S058 | OC | 0.38 | 27 | 18 | 65 | 3.20 | 60.40 | Y |
| S059 | OC | 0.63 | 27 | 17 | 44 | 1.81 | 59.95 | N |
| S060 | OC | 0.45 | 29 | 17 | 38 | 1.71 | 57.50 | Y |
| S061 | OC | 0.29 | 25 | 19 | 61 | 3.06 | 57.57 | Y |
| S062 | BC | 0.69 | 30 | 33 | 18 | 1.55 | 56.98 | Y |
| S063 | OC | 0.64 | 25 | 20 | 62 | 3.25 | 56.63 | Y |
| S064 | OC | 0.67 | 31 | 16 | 54 | 2.90 | 56.05 | Y |
| S065 | BC | 0.20 | 29 | 48 | 13 | 2.20 | 55.90 | N |
| S066 | OC | 0.55 | 27 | 23 | 43 | 2.39 | 55.96 | N |
| S067 | OC | 0.78 | 24 | 19 | 47 | 2.28 | 54.66 | N |
| S068 | OC | 0.46 | 22 | 24 | 38 | 1.90 | 54.55 | Y |
| S069 | OC | 0.51 | 28 | 18 | 57 | 3.16 | 54.02 | N |
| S070 | OC | 0.74 | 23 | 18 | 40 | 1.75 | 53.88 | Y |
| S071 | OC | 0.50 | 25 | 15 | 44 | 1.99 | 53.16 | N |
| S072 | OC | 0.33 | 30 | 17 | 43 | 2.43 | 52.34 | Y |
| S073 | OC | 0.47 | 15 | 20 | 69 | 3.34 | 52.23 | Y |
| S074 | OC | 0.50 | 25 | 19 | 35 | 1.84 | 50.48 | N |
| S075 | OC | 0.80 | 24 | 14 | 44 | 2.05 | 50.23 | N |
| S076 | OC | 0.36 | 19 | 16 | 65 | 3.29 | 49.01 | N |
| S077 | BC | 0.64 | 26 | 38 | 15 | 1.95 | 48.78 | Y |
| S078 | OC | 0.53 | 24 | 14 | 38 | 1.77 | 48.57 | Y |
| S079 | OC | 0.82 | 24 | 18 | 34 | 1.79 | 48.26 | N |
| S080 | BC | 0.56 | 23 | 35 | 18 | 1.82 | 47.79 | Y |
| S081 | OC | 0.56 | 25 | 17 | 32 | 1.72 | 47.34 | Y |
| S082 | OC | 0.88 | 23 | 19 | 32 | 1.74 | 47.03 | Y |
| S083 | OC | 0.77 | 11 | 20 | 74 | 3.74 | 47.03 | N |
| S084 | OC | 0.37 | 25 | 19 | 32 | 1.88 | 46.86 | N |
| S085 | OC | 0.74 | 17 | 19 | 61 | 3.28 | 46.16 | Y |
| S086 | OC | 0.73 | 25 | 16 | 38 | 2.14 | 45.83 | Y |
| S087 | OC | 0.74 | 27 | 14 | 37 | 2.08 | 45.76 | Y |
| S088 | OC | 0.76 | 24 | 14 | 34 | 1.72 | 45.34 | Y |
| S089 | BC | 0.33 | 26 | 31 | 13 | 1.61 | 45.05 | N |
| S090 | OC | 0.47 | 24 | 17 | 49 | 2.91 | 44.90 | Y |
| S091 | OC | 0.62 | 23 | 15 | 36 | 1.88 | 44.86 | N |
| S092 | OC | 0.32 | 24 | 19 | 45 | 2.79 | 44.76 | Y |
| S093 | OC | 0.62 | 23 | 12 | 37 | 1.82 | 43.79 | N |
| S094 | BC | 0.77 | 23 | 34 | 17 | 1.96 | 43.62 | Y |
| S095 | OC | 0.46 | 19 | 17 | 36 | 1.90 | 42.55 | N |
| S096 | OC | 0.37 | 21 | 18 | 51 | 3.12 | 41.64 | Y |
| S097 | OC | 0.83 | 23 | 12 | 34 | 1.77 | 41.57 | N |
| S098 | OC | 0.82 | 22 | 17 | 32 | 1.92 | 41.24 | N |
| S099 | OC | 0.66 | 19 | 17 | 49 | 2.86 | 40.67 | Y |
| S100 | OC | 0.24 | 24 | 16 | 36 | 2.30 | 40.35 | N |
| S101 | OC | 0.61 | 17 | 18 | 54 | 3.14 | 40.33 | Y |
| S102 | OC | 0.65 | 21 | 12 | 37 | 1.94 | 39.93 | N |
| S103 | OC | 0.86 | 22 | 16 | 29 | 1.77 | 39.57 | Y |
| S104 | OC | 0.66 | 22 | 20 | 46 | 3.15 | 39.18 | Y |
| S105 | BC | 0.44 | 23 | 26 | 17 | 1.76 | 38.72 | N |
| S106 | OC | 0.32 | 11 | 10 | 56 | 2.49 | 38.41 | Y |
| S107 | OC | 0.29 | 22 | 13 | 30 | 1.74 | 38.03 | N |
| S108 | OC | 0.79 | 23 | 20 | 37 | 2.72 | 37.84 | N |
| S109 | OC | 0.64 | 21 | 16 | 30 | 1.92 | 37.24 | N |
| S110 | OC | 0.70 | 19 | 16 | 29 | 1.75 | 36.88 | N |
| S111 | OC | 0.26 | 22 | 13 | 31 | 1.90 | 36.55 | N |
| S112 | OC | 0.56 | 21 | 15 | 27 | 1.72 | 36.34 | Y |
| S113 | OC | 0.54 | 13 | 19 | 42 | 2.45 | 36.03 | N |
| S114 | OC | 0.43 | 20 | 15 | 29 | 1.84 | 35.48 | N |
| S115 | OC | 0.71 | 21 | 12 | 32 | 1.92 | 35.24 | Y |
| S116 | BC | 0.48 | 18 | 55 | 18 | 3.62 | 34.89 | Y |
| S117 | OC | 0.31 | 22 | 14 | 30 | 2.05 | 34.23 | Y |
| S118 | OC | 0.79 | 16 | 19 | 29 | 1.93 | 34.09 | Y |
| S119 | OC | 0.65 | 23 | 13 | 25 | 1.81 | 32.95 | Y |
| S120 | BC | 0.43 | 17 | 38 | 21 | 2.78 | 32.91 | N |
| S121 | OC | 0.63 | 18 | 11 | 37 | 2.16 | 32.52 | N |
| S122 | OC | 0.44 | 17 | 6 | 51 | 2.72 | 31.84 | Y |
| S123 | OC | 0.53 | 16 | 14 | 38 | 2.34 | 31.73 | Y |
| S124 | OC | 0.68 | 15 | 14 | 32 | 1.89 | 31.71 | N |
| S125 | OC | 0.63 | 14 | 13 | 47 | 2.74 | 31.53 | N |
| S126 | OC | 0.21 | 17 | 9 | 42 | 2.37 | 31.27 | N |
| S127 | OC | 0.20 | 20 | 11 | 36 | 2.37 | 30.27 | Y |
| S128 | OC | 0.78 | 22 | 12 | 22 | 1.67 | 30.12 | N |
| S129 | OC | 0.52 | 18 | 13 | 26 | 1.80 | 29.10 | N |
| S130 | OC | 0.67 | 19 | 12 | 27 | 1.87 | 29.02 | N |
| S131 | OC | 0.46 | 16 | 17 | 26 | 1.94 | 28.93 | N |
| S132 | OC | 0.44 | 17 | 23 | 36 | 3.06 | 28.57 | N |
| S133 | OC | 0.67 | 18 | 17 | 42 | 3.16 | 28.02 | N |
| S134 | BC | 0.40 | 14 | 28 | 17 | 2.00 | 28.00 | N |
| S135 | OC | 0.70 | 9 | 17 | 53 | 3.35 | 27.08 | N |
| S136 | OC | 0.36 | 16 | 13 | 25 | 1.75 | 26.88 | N |
| S137 | OC | 0.27 | 19 | 14 | 22 | 1.87 | 26.02 | N |
| S138 | OC | 0.40 | 16 | 11 | 43 | 2.84 | 25.98 | N |
| S139 | OC | 0.56 | 16 | 15 | 24 | 1.89 | 25.71 | N |
| S140 | OC | 0.85 | 19 | 9 | 19 | 1.41 | 25.15 | N |
| S141 | OC | 0.60 | 19 | 9 | 25 | 1.87 | 24.02 | N |
| S142 | OC | 0.46 | 14 | 13 | 37 | 2.59 | 23.86 | N |
| S143 | OC | 0.23 | 18 | 10 | 38 | 2.02 | 34.68 | Y |
| S144 | OC | 0.30 | 11 | 51 | 5 | 2.79 | 23.76 | N |
| S145 | OC | 0.78 | 14 | 14 | 26 | 1.96 | 23.62 | N |
| S146 | OC | 0.40 | 15 | 7 | 26 | 1.59 | 23.36 | N |
| S147 | OC | 0.84 | 16 | 13 | 22 | 1.80 | 23.10 | N |
| S148 | BC | 0.27 | 14 | 24 | 16 | 1.99 | 23.16 | Y |
| S149 | BC | 0.41 | 8 | 61 | 15 | 3.99 | 22.16 | N |
| S150 | OC | 0.36 | 16 | 13 | 22 | 1.90 | 21.55 | N |
| S151 | OC | 0.88 | 17 | 9 | 37 | 2.73 | 20.69 | N |
| S152 | OC | 0.42 | 16 | 11 | 26 | 2.11 | 20.30 | N |
| S153 | OC | 0.50 | 16 | 16 | 43 | 3.55 | 19.98 | N |
| S154 | BC | 0.35 | 16 | 33 | 15 | 2.92 | 18.74 | N |
| S155 | BC | 0.48 | 14 | 26 | 11 | 2.15 | 17.68 | N |
| S156 | OC | 0.41 | 15 | 16 | 37 | 3.25 | 17.63 | N |
| S157 | OC | 0.67 | 16 | 12 | 19 | 1.96 | 16.62 | N |
| S158 | BC | 0.43 | 13 | 22 | 12 | 1.97 | 16.47 | N |
| S159 | OC | 0.47 | 12 | 13 | 34 | 2.83 | 15.14 | N |
| S160 | OC | 0.86 | 15 | 7 | 19 | 1.68 | 14.96 | N |
| S161 | BC | 0.38 | 14 | 24 | 8 | 2.10 | 13.45 | N |
| S162 | OC | 0.41 | 13 | 14 | 41 | 3.52 | 13.44 | N |
| S163 | OC | 0.76 | 16 | 12 | 14 | 1.86 | 13.17 | N |
| S164 | BC | 0.36 | 16 | 18 | 6 | 1.74 | 13.03 | N |
| S165 | OC | 0.88 | 14 | 7 | 19 | 1.75 | 12.88 | N |
| S166 | OC | 0.60 | 8 | 15 | 42 | 3.38 | 12.61 | N |
| S167 | OC | 0.78 | 5 | 14 | 40 | 3.15 | 10.18 | N |
| S168 | OC | 0.25 | 15 | 10 | 19 | 2.19 | 10.06 | N |
| S169 | OC | 0.54 | 15 | 5 | 23 | 2.13 | 9.99 | N |
| S170 | OC | 0.41 | 11 | 16 | 33 | 3.23 | 9.94 | N |
| S171 | OC | 0.21 | 11 | 9 | 28 | 2.47 | 9.72 | N |
| S172 | OC | 0.77 | 5 | 6 | 31 | 2.21 | 7.75 | N |
| S173 | BC | 0.46 | 11 | 16 | 9 | 1.85 | 7.33 | N |
| S174 | OC | 0.82 | 14 | 11 | 31 | 3.19 | 6.56 | N |
| S175 | BC | 0.20 | 13 | 12 | 19 | 2.43 | 6.34 | N |
| S176 | BC | 0.47 | 13 | 15 | 6 | 1.79 | 6.26 | N |
| S177 | OC | 0.36 | 9 | 12 | 18 | 2.15 | 5.68 | N |
| S178 | BC | 0.38 | 11 | 15 | 9 | 1.92 | 5.24 | N |
| S179 | OC | 0.71 | 6 | 9 | 41 | 3.41 | 3.15 | N |
| S180 | OC | 0.53 | 8 | 15 | 33 | 3.42 | 2.99 | N |
| S181 | OC | 0.37 | 8 | 11 | 36 | 3.40 | 2.30 | N |
| S182 | BC | 0.41 | 9 | 17 | 6 | 2.05 | 0.23 | N |
| S183 | OC | 0.64 | 7 | 12 | 28 | 3.03 | 0.04 | N |
| S184 | OC | 0.80 | 12 | 5 | 14 | 2.01 | -0.15 | N |
| S185 | OC | 0.63 | 12 | 6 | 15 | 2.17 | -0.63 | N |
| S186 | OC | 0.28 | 11 | 8 | 17 | 2.59 | -4.15 | N |
| S187 | BC | 0.32 | 7 | 16 | 4 | 2.04 | -4.62 | N |
| S188 | OC | 0.64 | 4 | 7 | 40 | 3.69 | -6.20 | N |
| S189 | BC | 0.60 | 6 | 11 | 7 | 1.98 | -6.69 | N |
| S190 | BC | 0.35 | 4 | 17 | 7 | 2.50 | -10.75 | N |
| S191 | OC | 0.35 | 6 | 4 | 8 | 1.93 | -11.92 | N |
| S192 | OC | 0.26 | 6 | 5 | 7 | 2.10 | -14.55 | Y |
| S193 | OC | 0.63 | 2 | 11 | 32 | 3.86 | -14.83 | N |
| S194 | OC | 0.77 | 3 | 3 | 4 | 2.10 | -22.55 | N |
| S195 | OC | 0.83 | 0 | 3 | 24 | 4.07 | -36.09 | N |

*Y means that BRCA1/2 is deficient, and N means that *BRCA1/2* is intact.

Supplemental Table 4 Comparison of the chromosome segmentation and copy number of GSA, PureCN, and ASCAT.

| GSA | | | | | PureCN | | | | ASCAT | | | | |
| --- | --- | --- | --- | --- | --- | --- | --- | --- | --- | --- | --- | --- | --- |
| chromosome | start | end | nA | nB | chromosome | start | end | Copy number | chromosome | start | end | nMajor | nMinor |
| 1 | 10443 | 3683916 | 5 | 5 | 1 | 10544 | 108910866 | 3 | 1 | 10444 | 255957 | 2 | 0 |
| 1 | 3683916 | 16797479 | 2 | 2 | 1 | 110246770 | 121350186 | 3 | 1 | 534242 | 1509825 | 14 | 8 |
| 1 | 16797480 | 22451377 | 3 | 3 | 1 | 121390914 | 142728945 | 6 | 1 | 1534614 | 3683916 | 7 | 6 |
| 1 | 22451378 | 74054262 | 2 | 2 | 1 | 142811022 | 228198520 | 3 | 1 | 3686315 | 5748792 | 3 | 3 |
| 1 | 74054263 | 78149691 | 2 | 3 | 1 | 228617059 | 231790931 | 3 | 1 | 5765147 | 6695123 | 6 | 6 |
| 1 | 78149692 | 120383827 | 2 | 2 | 1 | 231790932 | 249239442 | 2 | 1 | 6700095 | 16054700 | 4 | 3 |
| 1 | 120383828 | 150059069 | 2 | 3 | 2 | 10287 | 20845260 | 3 | 1 | 16063698 | 17331676 | 9 | 6 |
| 1 | 150059069 | 164109310 | 2 | 2 | 2 | 20870815 | 89560394 | 3 | 1 | 17342924 | 33851400 | 4 | 3 |
| 1 | 164109311 | 177669266 | 1 | 2 | 2 | 91757142 | 146858894 | 3 | 1 | 33858796 | 40768552 | 3 | 3 |
| 1 | 177669267 | 201016435 | 2 | 2 | 2 | 146885790 | 240957786 | 3 | 1 | 40769415 | 45297753 | 4 | 3 |
| 1 | 201016436 | 206694171 | 2 | 3 | 3 | 105785 | 97983265 | 3 | 1 | 45307506 | 51705962 | 3 | 2 |
| 1 | 206694171 | 249239610 | 2 | 2 | 3 | 97983266 | 129279827 | 3 | 1 | 51730054 | 55418976 | 3 | 3 |
| 2 | 12990 | 75590677 | 1 | 2 | 3 | 129331896 | 183907028 | 3 | 1 | 55429134 | 55538385 | 7 | 6 |
| 2 | 75590677 | 173916192 | 2 | 2 | 3 | 184312097 | 197860922 | 2 | 1 | 55541174 | 74379335 | 3 | 2 |
| 2 | 173916193 | 177912470 | 2 | 3 | 3 | 184312097 | 197860922 | 2 | 1 | 74388971 | 77828321 | 4 | 2 |
| 2 | 177912471 | 241066575 | 2 | 2 | 4 | 10798 | 9071586 | 3 | 1 | 77840200 | 85036239 | 2 | 2 |
| 2 | 241066575 | 243184927 | 4 | 4 | 4 | 9093573 | 15004879 | 4 | 1 | 85040069 | 85746754 | 5 | 3 |
| 3 | 105785 | 22966271 | 1 | 2 | 4 | 15009222 | 69382795 | 3 | 1 | 85768379 | 109650386 | 3 | 2 |
| 3 | 22966271 | 25149971 | 1 | 3 | 4 | 70223753 | 190939656 | 3 | 1 | 109656420 | 110168888 | 6 | 4 |
| 3 | 25149971 | 31998548 | 1 | 2 | 5 | 2909107 | 42674475 | 3 | 1 | 110169957 | 110254678 | 8 | 5 |
| 3 | 31998549 | 35044116 | 1 | 3 | 5 | 42674476 | 100669673 | 3 | 1 | 110268828 | 113123734 | 3 | 3 |
| 3 | 35044117 | 64619211 | 1 | 2 | 5 | 100669674 | 180719384 | 3 | 1 | 113140583 | 113290718 | 6 | 4 |
| 3 | 64619212 | 67527113 | 1 | 3 | 6 | 183913 | 29855898 | 3 | 1 | 113310636 | 120256713 | 3 | 2 |
| 3 | 67527114 | 89232164 | 1 | 2 | 6 | 34078428 | 125368280 | 3 | 1 | 120267544 | 121449630 | 5 | 1 |
| 3 | 89232165 | 164773057 | 2 | 2 | 6 | 125368281 | 166718428 | 3 | 1 | 142540154 | 143505099 | 3 | 1 |
| 3 | 164773057 | 197861082 | 2 | 3 | 6 | 166736361 | 170978749 | 3 | 1 | 143509346 | 156206115 | 4 | 3 |
| 4 | 11870 | 3789961 | 2 | 2 | 7 | 67812 | 54497365 | 3 | 1 | 156213912 | 156909303 | 5 | 5 |
| 4 | 3789961 | 190907201 | 1 | 1 | 7 | 57698751 | 61551813 | 3 | 1 | 156914070 | 164182915 | 3 | 2 |
| 5 | 11788 | 2413820 | 2 | 6 | 7 | 61751405 | 76144500 | 3 | 1 | 164205963 | 177698891 | 2 | 1 |
| 5 | 2413820 | 45414358 | 1 | 3 | 7 | 76198922 | 84604610 | 3 | 1 | 177707062 | 185487243 | 3 | 2 |
| 5 | 50329524 | 59284115 | 1 | 1 | 7 | 88213051 | 118686043 | 3 | 1 | 185500360 | 197407942 | 2 | 2 |
| 5 | 59284116 | 64758380 | 1 | 2 | 7 | 118884560 | 141786093 | 3 | 1 | 197414077 | 200838925 | 3 | 3 |
| 5 | 64758381 | 100730500 | 1 | 1 | 7 | 143036338 | 147182357 | 3 | 1 | 200842104 | 200956288 | 7 | 6 |
| 5 | 100730501 | 137802107 | 1 | 2 | 7 | 147182358 | 159128528 | 3 | 1 | 200958111 | 201116018 | 9 | 8 |
| 5 | 137802108 | 140959743 | 2 | 2 | 8 | 10470 | 3172400 | 4 | 1 | 201122947 | 201358297 | 6 | 5 |
| 5 | 140959744 | 180719407 | 1 | 2 | 8 | 3190640 | 7222674 | 4 | 1 | 201359717 | 204397427 | 4 | 3 |
| 6 | 184125 | 122114448 | 1 | 1 | 8 | 7868986 | 11878174 | 4 | 1 | 204403659 | 205499698 | 7 | 4 |
| 6 | 122114448 | 125597537 | 1 | 2 | 8 | 12575672 | 37190826 | 4 | 1 | 205505474 | 225927179 | 2 | 2 |
| 6 | 125597537 | 171049459 | 0 | 1 | 8 | 39390215 | 115234266 | 3 | 1 | 225955919 | 227913886 | 3 | 2 |
| 7 | 34918 | 2758932 | 2 | 4 | 8 | 115240752 | 129073961 | 2 | 1 | 227920061 | 228612838 | 6 | 5 |
| 7 | 2758932 | 88774612 | 1 | 2 | 8 | 129077999 | 140789846 | 2 | 1 | 228617063 | 228699477 | 3 | 2 |
| 7 | 88774613 | 91497419 | 1 | 3 | 8 | 142021694 | 146301295 | 2 | 1 | 228714745 | 241728046 | 2 | 2 |
| 7 | 91497420 | 147891107 | 1 | 2 | 9 | 10308 | 20603357 | 3 | 1 | 241755341 | 242109146 | 5 | 2 |
| 7 | 147891108 | 159127624 | 1 | 3 | 9 | 20626651 | 28761500 | 4 | 1 | 242122950 | 249239246 | 2 | 2 |
| 8 | 11892 | 31255894 | 1 | 2 | 9 | 28761501 | 95708348 | 3 | 2 | 10437 | 3605468 | 3 | 2 |
| 8 | 31255895 | 36469972 | 1 | 1 | 9 | 96091676 | 127201626 | 3 | 2 | 3610367 | 29183332 | 3 | 1 |
| 8 | 36469973 | 39694832 | 1 | 2 | 9 | 129377847 | 141069049 | 3 | 2 | 29205451 | 29338016 | 6 | 3 |
| 8 | 39694833 | 139768085 | 2 | 2 | 10 | 2324863 | 51874242 | 3 | 2 | 29347233 | 43927509 | 2 | 1 |
| 8 | 139768085 | 142199400 | 3 | 4 | 10 | 51911792 | 69482430 | 3 | 2 | 43931321 | 44099433 | 8 | 2 |
| 8 | 142199400 | 146300855 | 5 | 5 | 10 | 69528410 | 73782236 | 3 | 2 | 44101538 | 44502788 | 9 | 2 |
| 9 | 10345 | 30951193 | 1 | 1 | 10 | 73782237 | 79370711 | 3 | 2 | 44504877 | 75590677 | 2 | 1 |
| 9 | 30951194 | 125570106 | 1 | 2 | 10 | 79403929 | 103501765 | 3 | 2 | 75612112 | 95847037 | 3 | 2 |
| 9 | 125570107 | 129864419 | 2 | 2 | 10 | 103501766 | 133218174 | 3 | 2 | 95862265 | 98413053 | 4 | 3 |
| 9 | 129864420 | 133967240 | 2 | 4 | 11 | 193151 | 3315255 | 3 | 2 | 98419726 | 113817566 | 3 | 2 |
| 9 | 133967240 | 136119809 | 1 | 2 | 11 | 3362845 | 8657322 | 4 | 2 | 113819614 | 114035194 | 4 | 4 |
| 9 | 136119809 | 141034077 | 3 | 5 | 11 | 8707150 | 15267597 | 3 | 2 | 114044377 | 120436681 | 3 | 2 |
| 10 | 135708 | 2989837 | 1 | 1 | 11 | 15267598 | 64012335 | 3 | 2 | 120438515 | 121854242 | 4 | 3 |
| 10 | 2989837 | 7063049 | 1 | 5 | 11 | 70605729 | 77069740 | 3 | 2 | 121873228 | 126322442 | 2 | 2 |
| 10 | 7063050 | 38181470 | 1 | 2 | 11 | 77111279 | 124249831 | 3 | 2 | 126335702 | 130860632 | 3 | 2 |
| 10 | 42546687 | 47006215 | 1 | 3 | 11 | 124249832 | 134946332 | 3 | 2 | 130870631 | 132302165 | 4 | 4 |
| 10 | 47006216 | 72990780 | 1 | 2 | 12 | 193817 | 9619771 | 3 | 2 | 132312325 | 173884290 | 2 | 2 |
| 10 | 72990781 | 77392753 | 2 | 5 | 12 | 9745645 | 14705349 | 3 | 2 | 173913258 | 178417225 | 3 | 3 |
| 10 | 77392754 | 103535470 | 1 | 3 | 12 | 14748352 | 24215922 | 3 | 2 | 178430747 | 216863221 | 2 | 2 |
| 10 | 103535471 | 132854084 | 2 | 3 | 12 | 26581301 | 54410000 | 3 | 2 | 216878215 | 218674353 | 4 | 3 |
| 10 | 132854084 | 135523863 | 5 | 7 | 12 | 54756526 | 133839682 | 3 | 2 | 218675144 | 220537405 | 4 | 4 |
| 11 | 193096 | 3254265 | 0 | 5 | 13 | 19365400 | 115108995 | 3 | 2 | 220543947 | 233272520 | 3 | 2 |
| 11 | 3254265 | 15287427 | 0 | 2 | 14 | 20369396 | 41602932 | 3 | 2 | 233273244 | 233410503 | 8 | 7 |
| 11 | 15287428 | 63668557 | 1 | 2 | 14 | 41676644 | 73181555 | 3 | 2 | 233415520 | 238166147 | 3 | 3 |
| 11 | 63668557 | 65906059 | 4 | 4 | 14 | 74040334 | 82491999 | 3 | 2 | 238172570 | 238503559 | 6 | 4 |
| 11 | 65906060 | 71329189 | 2 | 4 | 14 | 82503803 | 106412532 | 3 | 2 | 238509769 | 243184927 | 5 | 4 |
| 11 | 71329189 | 117054709 | 1 | 2 | 15 | 24752517 | 30432873 | 3 | 3 | 105785 | 9798006 | 2 | 1 |
| 11 | 117054709 | 120387112 | 2 | 3 | 15 | 30889846 | 93588316 | 3 | 3 | 9798767 | 9991369 | 4 | 3 |
| 11 | 120387112 | 134946381 | 1 | 2 | 15 | 93616927 | 102431165 | 3 | 3 | 10015437 | 12854616 | 2 | 1 |
| 12 | 193817 | 44325870 | 1 | 2 | 16 | 2998646 | 32096230 | 3 | 3 | 12856856 | 13726888 | 4 | 2 |
| 12 | 44325871 | 52856128 | 1 | 3 | 16 | 33839127 | 55778841 | 3 | 3 | 13734613 | 23503506 | 2 | 1 |
| 12 | 52856129 | 69518950 | 1 | 2 | 16 | 55832532 | 90171440 | 3 | 3 | 23511338 | 24830136 | 3 | 1 |
| 12 | 69518950 | 72779820 | 2 | 3 | 17 | 301 | 15039064 | 3 | 3 | 24838218 | 31981493 | 2 | 1 |
| 12 | 72779820 | 130897316 | 1 | 2 | 17 | 18221011 | 21982191 | 3 | 3 | 32001641 | 35035432 | 4 | 1 |
| 12 | 130897316 | 133840496 | 2 | 3 | 17 | 21982192 | 26633385 | 3 | 3 | 35045130 | 38034821 | 2 | 1 |
| 13 | 19149491 | 35918811 | 1 | 2 | 17 | 27816679 | 36211682 | 3 | 3 | 38035508 | 38793940 | 3 | 2 |
| 13 | 35918812 | 38039200 | 1 | 3 | 17 | 38140545 | 45490251 | 3 | 3 | 38802251 | 48384721 | 2 | 1 |
| 13 | 38039200 | 97966870 | 1 | 2 | 17 | 45490252 | 81050912 | 3 | 3 | 48393556 | 48678011 | 5 | 3 |
| 13 | 97966871 | 100749907 | 1 | 3 | 18 | 10718 | 77674628 | 4 | 3 | 48678971 | 48723302 | 7 | 4 |
| 13 | 100749908 | 115104231 | 1 | 2 | 19 | 844021 | 14924875 | 3 | 3 | 48727112 | 49928691 | 3 | 1 |
| 14 | 19066305 | 104518453 | 1 | 2 | 19 | 24487371 | 38793310 | 2 | 3 | 49936102 | 50645793 | 7 | 3 |
| 14 | 104518453 | 107289541 | 2 | 4 | 19 | 41104875 | 57804362 | 3 | 3 | 50647888 | 51697493 | 2 | 1 |
| 15 | 20119124 | 95416418 | 1 | 1 | 20 | 76963 | 9806342 | 3 | 3 | 51737965 | 52512318 | 4 | 2 |
| 15 | 95416419 | 102520957 | 1 | 2 | 20 | 9836329 | 26072253 | 3 | 3 | 52521921 | 52584424 | 7 | 4 |
| 16 | 103517 | 5196763 | 2 | 2 | 20 | 26278201 | 29652157 | 4 | 3 | 52584715 | 53277645 | 3 | 2 |
| 16 | 5196763 | 28336882 | 1 | 1 | 20 | 29809688 | 33585427 | 3 | 3 | 53293157 | 65306638 | 2 | 1 |
| 16 | 28336882 | 33965813 | 2 | 2 | 20 | 33591112 | 38747958 | 3 | 3 | 65311745 | 67425983 | 4 | 1 |
| 16 | 33965813 | 69525862 | 1 | 1 | 20 | 39998485 | 52169749 | 2 | 3 | 67431071 | 88916705 | 2 | 1 |
| 16 | 69525862 | 72832651 | 0 | 2 | 20 | 55892520 | 62916447 | 2 | 3 | 88926656 | 123005476 | 3 | 2 |
| 16 | 72832651 | 87518783 | 1 | 1 | 21 | 9527025 | 14572806 | 4 | 3 | 123010363 | 123135244 | 5 | 5 |
| 16 | 87518783 | 90230101 | 2 | 2 | 21 | 14578435 | 40683477 | 3 | 3 | 123152897 | 123419733 | 5 | 4 |
| 17 | 2037 | 8851027 | 0 | 3 | 21 | 40712373 | 48118698 | 3 | 3 | 123440967 | 123457711 | 8 | 7 |
| 17 | 8851027 | 33194247 | 0 | 2 | 22 | 24681539 | 30638282 | 3 | 3 | 123479778 | 125962089 | 3 | 2 |
| 17 | 33194248 | 45664593 | 0 | 1 | 22 | 30802434 | 51243277 | 3 | 3 | 125975564 | 129349136 | 4 | 4 |
| 17 | 45664593 | 48761099 | 1 | 3 |  |  |  |  | 3 | 129364254 | 147066494 | 3 | 2 |
| 17 | 48761099 | 69119481 | 1 | 2 |  |  |  |  | 3 | 147076030 | 147599269 | 4 | 4 |
| 17 | 69119481 | 78765219 | 2 | 4 |  |  |  |  | 3 | 147623909 | 164764719 | 2 | 2 |
| 17 | 78765219 | 81194987 | 2 | 5 |  |  |  |  | 3 | 164773057 | 175513789 | 4 | 3 |
| 18 | 10813 | 78017073 | 0 | 2 |  |  |  |  | 3 | 175520406 | 183907032 | 3 | 3 |
| 19 | 282747 | 4818373 | 0 | 2 |  |  |  |  | 3 | 183914759 | 184311940 | 7 | 7 |
| 19 | 4818373 | 14231330 | 2 | 6 |  |  |  |  | 3 | 184320458 | 184485565 | 5 | 4 |
| 19 | 14231330 | 17296679 | 1 | 2 |  |  |  |  | 3 | 184492965 | 193798407 | 3 | 3 |
| 19 | 17296679 | 20184274 | 2 | 4 |  |  |  |  | 3 | 193807621 | 197861082 | 5 | 4 |
| 19 | 20184274 | 22482889 | 1 | 4 |  |  |  |  | 4 | 10798 | 515489 | 1 | 1 |
| 19 | 22482890 | 38604588 | 1 | 1 |  |  |  |  | 4 | 515844 | 2074700 | 3 | 2 |
| 19 | 38604588 | 41284394 | 2 | 2 |  |  |  |  | 4 | 2087202 | 2307960 | 1 | 1 |
| 19 | 41284394 | 48785642 | 1 | 4 |  |  |  |  | 4 | 2330889 | 3789961 | 2 | 1 |
| 19 | 48785642 | 56767042 | 2 | 4 |  |  |  |  | 4 | 3798866 | 6525332 | 1 | 1 |
| 19 | 56767042 | 59118886 | 1 | 2 |  |  |  |  | 4 | 6532811 | 9022171 | 2 | 1 |
| 20 | 76962 | 25041159 | 2 | 2 |  |  |  |  | 4 | 9032235 | 190939879 | 1 | 0 |
| 20 | 30273552 | 49774398 | 2 | 2 |  |  |  |  | 5 | 12114 | 14104 | 9 | 0 |
| 20 | 49774399 | 53291424 | 2 | 4 |  |  |  |  | 5 | 14757 | 49851 | 3 | 1 |
| 20 | 53291425 | 56433748 | 2 | 3 |  |  |  |  | 5 | 50708 | 1889840 | 8 | 3 |
| 20 | 56433749 | 60180124 | 2 | 2 |  |  |  |  | 5 | 1902586 | 42779736 | 3 | 1 |
| 20 | 60180124 | 62958710 | 4 | 4 |  |  |  |  | 5 | 42782492 | 43855368 | 4 | 3 |
| 21 | 9527088 | 32935103 | 2 | 2 |  |  |  |  | 5 | 43927178 | 45387846 | 3 | 1 |
| 21 | 32935103 | 35506430 | 2 | 3 |  |  |  |  | 5 | 45406321 | 49505838 | 2 | 1 |
| 21 | 35506430 | 43852421 | 2 | 2 |  |  |  |  | 5 | 49522346 | 59277134 | 1 | 0 |
| 21 | 43852422 | 48104004 | 4 | 4 |  |  |  |  | 5 | 59286205 | 64693317 | 2 | 1 |
| 22 | 16870802 | 24655659 | 0 | 3 |  |  |  |  | 5 | 64725524 | 79238533 | 1 | 0 |
| 22 | 24655659 | 28604583 | 0 | 2 |  |  |  |  | 5 | 79252182 | 79617357 | 2 | 1 |
| 22 | 28604583 | 31438928 | 0 | 4 |  |  |  |  | 5 | 79621016 | 94259610 | 1 | 0 |
| 22 | 31438928 | 51183255 | 0 | 2 |  |  |  |  | 5 | 94278129 | 94528491 | 2 | 1 |
|  |  |  |  |  |  |  |  |  | 5 | 94550971 | 100753611 | 1 | 0 |
|  |  |  |  |  |  |  |  |  | 5 | 100765147 | 138487483 | 2 | 1 |
|  |  |  |  |  |  |  |  |  | 5 | 138494690 | 138705812 | 2 | 2 |
|  |  |  |  |  |  |  |  |  | 5 | 138713959 | 141335241 | 4 | 3 |
|  |  |  |  |  |  |  |  |  | 5 | 141336264 | 148152364 | 2 | 1 |
|  |  |  |  |  |  |  |  |  | 5 | 148164206 | 148581063 | 6 | 2 |
|  |  |  |  |  |  |  |  |  | 5 | 148610276 | 149431259 | 3 | 1 |
|  |  |  |  |  |  |  |  |  | 5 | 149433596 | 150158164 | 4 | 2 |
|  |  |  |  |  |  |  |  |  | 5 | 150164243 | 167653394 | 2 | 1 |
|  |  |  |  |  |  |  |  |  | 5 | 167674363 | 168634740 | 6 | 2 |
|  |  |  |  |  |  |  |  |  | 5 | 168671698 | 175956516 | 2 | 1 |
|  |  |  |  |  |  |  |  |  | 5 | 175956516 | 176148768 | 6 | 3 |
|  |  |  |  |  |  |  |  |  | 5 | 176165086 | 180719407 | 4 | 2 |
|  |  |  |  |  |  |  |  |  | 6 | 184125 | 25918688 | 1 | 0 |
|  |  |  |  |  |  |  |  |  | 6 | 25944103 | 33272712 | 1 | 1 |
|  |  |  |  |  |  |  |  |  | 6 | 33280505 | 34110254 | 2 | 1 |
|  |  |  |  |  |  |  |  |  | 6 | 34124370 | 44691371 | 1 | 1 |
|  |  |  |  |  |  |  |  |  | 6 | 44698708 | 90565234 | 1 | 0 |
|  |  |  |  |  |  |  |  |  | 6 | 90572196 | 91204213 | 2 | 1 |
|  |  |  |  |  |  |  |  |  | 6 | 91225132 | 98280013 | 1 | 0 |
|  |  |  |  |  |  |  |  |  | 6 | 98300239 | 98749236 | 2 | 1 |
|  |  |  |  |  |  |  |  |  | 6 | 98771028 | 122605510 | 1 | 0 |
|  |  |  |  |  |  |  |  |  | 6 | 122617158 | 124201918 | 2 | 1 |
|  |  |  |  |  |  |  |  |  | 6 | 124204860 | 125572968 | 4 | 1 |
|  |  |  |  |  |  |  |  |  | 6 | 125597537 | 166554170 | 0 | 0 |
|  |  |  |  |  |  |  |  |  | 6 | 166572045 | 171050287 | 1 | 0 |
|  |  |  |  |  |  |  |  |  | 7 | 16165 | 2458380 | 5 | 2 |
|  |  |  |  |  |  |  |  |  | 7 | 2472707 | 3117838 | 7 | 3 |
|  |  |  |  |  |  |  |  |  | 7 | 3127215 | 5239360 | 3 | 1 |
|  |  |  |  |  |  |  |  |  | 7 | 5254227 | 5686704 | 6 | 3 |
|  |  |  |  |  |  |  |  |  | 7 | 5696125 | 6870635 | 3 | 1 |
|  |  |  |  |  |  |  |  |  | 7 | 6878096 | 43846603 | 2 | 1 |
|  |  |  |  |  |  |  |  |  | 7 | 43851539 | 45257233 | 4 | 2 |
|  |  |  |  |  |  |  |  |  | 7 | 45273264 | 54133053 | 2 | 1 |
|  |  |  |  |  |  |  |  |  | 7 | 54142421 | 54689570 | 2 | 2 |
|  |  |  |  |  |  |  |  |  | 7 | 54711985 | 72286028 | 2 | 1 |
|  |  |  |  |  |  |  |  |  | 7 | 72300323 | 72891754 | 3 | 2 |
|  |  |  |  |  |  |  |  |  | 7 | 72898228 | 72934078 | 3 | 0 |
|  |  |  |  |  |  |  |  |  | 7 | 72944370 | 76162076 | 4 | 2 |
|  |  |  |  |  |  |  |  |  | 7 | 76173536 | 89563399 | 2 | 1 |
|  |  |  |  |  |  |  |  |  | 7 | 89574638 | 91443183 | 4 | 1 |
|  |  |  |  |  |  |  |  |  | 7 | 91464218 | 98990416 | 2 | 1 |
|  |  |  |  |  |  |  |  |  | 7 | 99017733 | 102312228 | 3 | 2 |
|  |  |  |  |  |  |  |  |  | 7 | 102332448 | 147821399 | 2 | 1 |
|  |  |  |  |  |  |  |  |  | 7 | 147839308 | 149321637 | 4 | 2 |
|  |  |  |  |  |  |  |  |  | 7 | 149334780 | 149523584 | 11 | 4 |
|  |  |  |  |  |  |  |  |  | 7 | 149528146 | 149576819 | 10 | 4 |
|  |  |  |  |  |  |  |  |  | 7 | 149589663 | 159128605 | 5 | 2 |
|  |  |  |  |  |  |  |  |  | 8 | 10470 | 2226147 | 3 | 1 |
|  |  |  |  |  |  |  |  |  | 8 | 2239128 | 10438796 | 2 | 1 |
|  |  |  |  |  |  |  |  |  | 8 | 10448836 | 10649192 | 5 | 3 |
|  |  |  |  |  |  |  |  |  | 8 | 10656437 | 16059915 | 2 | 1 |
|  |  |  |  |  |  |  |  |  | 8 | 16066975 | 17581155 | 3 | 1 |
|  |  |  |  |  |  |  |  |  | 8 | 17581700 | 19221650 | 2 | 1 |
|  |  |  |  |  |  |  |  |  | 8 | 19237079 | 20110641 | 3 | 2 |
|  |  |  |  |  |  |  |  |  | 8 | 20120297 | 21862551 | 2 | 1 |
|  |  |  |  |  |  |  |  |  | 8 | 21862551 | 23290439 | 4 | 2 |
|  |  |  |  |  |  |  |  |  | 8 | 23291833 | 25460335 | 2 | 1 |
|  |  |  |  |  |  |  |  |  | 8 | 25481266 | 26988103 | 1 | 0 |
|  |  |  |  |  |  |  |  |  | 8 | 27001935 | 31352513 | 2 | 1 |
|  |  |  |  |  |  |  |  |  | 8 | 31388393 | 35916718 | 1 | 0 |
|  |  |  |  |  |  |  |  |  | 8 | 35922774 | 39685594 | 2 | 1 |
|  |  |  |  |  |  |  |  |  | 8 | 39694860 | 43763025 | 3 | 2 |
|  |  |  |  |  |  |  |  |  | 8 | 43776238 | 47346329 | 4 | 4 |
|  |  |  |  |  |  |  |  |  | 8 | 47357126 | 84030205 | 2 | 2 |
|  |  |  |  |  |  |  |  |  | 8 | 84038956 | 123445622 | 3 | 2 |
|  |  |  |  |  |  |  |  |  | 8 | 123465838 | 140611843 | 3 | 3 |
|  |  |  |  |  |  |  |  |  | 8 | 140630990 | 142129925 | 7 | 4 |
|  |  |  |  |  |  |  |  |  | 8 | 142138860 | 145921475 | 9 | 8 |
|  |  |  |  |  |  |  |  |  | 8 | 145933692 | 146301381 | 4 | 3 |
|  |  |  |  |  |  |  |  |  | 9 | 10345 | 57539 | 2 | 1 |
|  |  |  |  |  |  |  |  |  | 9 | 58033 | 31146953 | 1 | 0 |
|  |  |  |  |  |  |  |  |  | 9 | 31164389 | 95826971 | 2 | 1 |
|  |  |  |  |  |  |  |  |  | 9 | 95840344 | 96060759 | 6 | 3 |
|  |  |  |  |  |  |  |  |  | 9 | 96072783 | 116741937 | 2 | 1 |
|  |  |  |  |  |  |  |  |  | 9 | 116751005 | 117195760 | 5 | 3 |
|  |  |  |  |  |  |  |  |  | 9 | 117203632 | 125840164 | 2 | 1 |
|  |  |  |  |  |  |  |  |  | 9 | 125848900 | 129551392 | 3 | 3 |
|  |  |  |  |  |  |  |  |  | 9 | 129558047 | 130210946 | 3 | 1 |
|  |  |  |  |  |  |  |  |  | 9 | 130213298 | 134530107 | 4 | 2 |
|  |  |  |  |  |  |  |  |  | 9 | 134541216 | 135841425 | 2 | 1 |
|  |  |  |  |  |  |  |  |  | 9 | 135861993 | 141069049 | 5 | 3 |
|  |  |  |  |  |  |  |  |  | 10 | 135708 | 1782394 | 1 | 1 |
|  |  |  |  |  |  |  |  |  | 10 | 1795194 | 3009351 | 1 | 0 |
|  |  |  |  |  |  |  |  |  | 10 | 3022248 | 3188071 | 15 | 4 |
|  |  |  |  |  |  |  |  |  | 10 | 3189380 | 6881554 | 8 | 2 |
|  |  |  |  |  |  |  |  |  | 10 | 6900363 | 38958568 | 2 | 1 |
|  |  |  |  |  |  |  |  |  | 10 | 38968721 | 42517060 | 3 | 1 |
|  |  |  |  |  |  |  |  |  | 10 | 42607765 | 47738963 | 5 | 2 |
|  |  |  |  |  |  |  |  |  | 10 | 47898043 | 71119003 | 2 | 1 |
|  |  |  |  |  |  |  |  |  | 10 | 71129194 | 73439101 | 3 | 2 |
|  |  |  |  |  |  |  |  |  | 10 | 73454637 | 73579217 | 6 | 3 |
|  |  |  |  |  |  |  |  |  | 10 | 73581560 | 73782236 | 3 | 1 |
|  |  |  |  |  |  |  |  |  | 10 | 73790819 | 77359185 | 7 | 2 |
|  |  |  |  |  |  |  |  |  | 10 | 77370319 | 80898955 | 4 | 1 |
|  |  |  |  |  |  |  |  |  | 10 | 80912499 | 81198212 | 7 | 3 |
|  |  |  |  |  |  |  |  |  | 10 | 81212704 | 92791385 | 4 | 1 |
|  |  |  |  |  |  |  |  |  | 10 | 92806230 | 96163034 | 6 | 2 |
|  |  |  |  |  |  |  |  |  | 10 | 96176376 | 103501914 | 4 | 1 |
|  |  |  |  |  |  |  |  |  | 10 | 103510619 | 110903958 | 4 | 4 |
|  |  |  |  |  |  |  |  |  | 10 | 110913556 | 111946947 | 4 | 1 |
|  |  |  |  |  |  |  |  |  | 10 | 111956363 | 133482891 | 4 | 4 |
|  |  |  |  |  |  |  |  |  | 10 | 133503598 | 135279810 | 10 | 9 |
|  |  |  |  |  |  |  |  |  | 10 | 135280765 | 135523863 | 7 | 2 |
|  |  |  |  |  |  |  |  |  | 11 | 193096 | 279786 | 3 | 1 |
|  |  |  |  |  |  |  |  |  | 11 | 280216 | 1961869 | 5 | 0 |
|  |  |  |  |  |  |  |  |  | 11 | 1971977 | 2073490 | 9 | 0 |
|  |  |  |  |  |  |  |  |  | 11 | 2094852 | 2436733 | 8 | 0 |
|  |  |  |  |  |  |  |  |  | 11 | 2439621 | 2441646 | 9 | 0 |
|  |  |  |  |  |  |  |  |  | 11 | 2443938 | 3254265 | 4 | 0 |
|  |  |  |  |  |  |  |  |  | 11 | 3260222 | 15282156 | 2 | 0 |
|  |  |  |  |  |  |  |  |  | 11 | 15301762 | 60504476 | 2 | 1 |
|  |  |  |  |  |  |  |  |  | 11 | 60508999 | 64039162 | 3 | 2 |
|  |  |  |  |  |  |  |  |  | 11 | 64053595 | 65617110 | 7 | 6 |
|  |  |  |  |  |  |  |  |  | 11 | 65623524 | 70517927 | 4 | 2 |
|  |  |  |  |  |  |  |  |  | 11 | 70523008 | 75852448 | 3 | 1 |
|  |  |  |  |  |  |  |  |  | 11 | 75860179 | 76924874 | 4 | 2 |
|  |  |  |  |  |  |  |  |  | 11 | 76928149 | 77814045 | 3 | 1 |
|  |  |  |  |  |  |  |  |  | 11 | 77815188 | 102587062 | 2 | 1 |
|  |  |  |  |  |  |  |  |  | 11 | 102593248 | 102870940 | 2 | 2 |
|  |  |  |  |  |  |  |  |  | 11 | 102884896 | 116634173 | 2 | 1 |
|  |  |  |  |  |  |  |  |  | 11 | 116643610 | 118472349 | 3 | 2 |
|  |  |  |  |  |  |  |  |  | 11 | 118478407 | 118604837 | 5 | 3 |
|  |  |  |  |  |  |  |  |  | 11 | 118626511 | 118757219 | 3 | 2 |
|  |  |  |  |  |  |  |  |  | 11 | 118757219 | 120151774 | 6 | 3 |
|  |  |  |  |  |  |  |  |  | 11 | 120161671 | 125714088 | 2 | 1 |
|  |  |  |  |  |  |  |  |  | 11 | 125747537 | 125986274 | 3 | 3 |
|  |  |  |  |  |  |  |  |  | 11 | 125998594 | 126232422 | 4 | 4 |
|  |  |  |  |  |  |  |  |  | 11 | 126242546 | 126404639 | 6 | 5 |
|  |  |  |  |  |  |  |  |  | 11 | 126411459 | 134946318 | 2 | 1 |
|  |  |  |  |  |  |  |  |  | 12 | 193817 | 907310 | 3 | 2 |
|  |  |  |  |  |  |  |  |  | 12 | 928911 | 6422272 | 2 | 1 |
|  |  |  |  |  |  |  |  |  | 12 | 6422988 | 7287914 | 4 | 2 |
|  |  |  |  |  |  |  |  |  | 12 | 7290200 | 44604028 | 2 | 1 |
|  |  |  |  |  |  |  |  |  | 12 | 44613070 | 48920006 | 4 | 1 |
|  |  |  |  |  |  |  |  |  | 12 | 48925862 | 50480232 | 3 | 1 |
|  |  |  |  |  |  |  |  |  | 12 | 50480314 | 52204246 | 2 | 1 |
|  |  |  |  |  |  |  |  |  | 12 | 52215102 | 53822992 | 4 | 2 |
|  |  |  |  |  |  |  |  |  | 12 | 53825325 | 57559809 | 2 | 1 |
|  |  |  |  |  |  |  |  |  | 12 | 57567180 | 57591458 | 7 | 4 |
|  |  |  |  |  |  |  |  |  | 12 | 57591458 | 57637593 | 6 | 3 |
|  |  |  |  |  |  |  |  |  | 12 | 57640620 | 69654187 | 2 | 1 |
|  |  |  |  |  |  |  |  |  | 12 | 69667075 | 70679929 | 3 | 3 |
|  |  |  |  |  |  |  |  |  | 12 | 70690194 | 72203204 | 3 | 1 |
|  |  |  |  |  |  |  |  |  | 12 | 72226611 | 113369644 | 2 | 1 |
|  |  |  |  |  |  |  |  |  | 12 | 113376452 | 113996719 | 4 | 4 |
|  |  |  |  |  |  |  |  |  | 12 | 114005404 | 120443494 | 2 | 1 |
|  |  |  |  |  |  |  |  |  | 12 | 120455954 | 121202362 | 6 | 2 |
|  |  |  |  |  |  |  |  |  | 12 | 121202952 | 122692820 | 3 | 2 |
|  |  |  |  |  |  |  |  |  | 12 | 122701001 | 124777480 | 1 | 1 |
|  |  |  |  |  |  |  |  |  | 12 | 124787539 | 125621419 | 3 | 2 |
|  |  |  |  |  |  |  |  |  | 12 | 125626535 | 131455442 | 2 | 1 |
|  |  |  |  |  |  |  |  |  | 12 | 131462375 | 132505406 | 4 | 2 |
|  |  |  |  |  |  |  |  |  | 12 | 132508389 | 133464490 | 7 | 7 |
|  |  |  |  |  |  |  |  |  | 12 | 133488904 | 133841396 | 3 | 2 |
|  |  |  |  |  |  |  |  |  | 13 | 19110523 | 35933714 | 2 | 1 |
|  |  |  |  |  |  |  |  |  | 13 | 35955344 | 37915301 | 4 | 1 |
|  |  |  |  |  |  |  |  |  | 13 | 37925269 | 98218375 | 2 | 1 |
|  |  |  |  |  |  |  |  |  | 13 | 98234789 | 100804923 | 4 | 2 |
|  |  |  |  |  |  |  |  |  | 13 | 100807376 | 110759711 | 2 | 1 |
|  |  |  |  |  |  |  |  |  | 13 | 110770593 | 113411956 | 3 | 2 |
|  |  |  |  |  |  |  |  |  | 13 | 113421845 | 115108995 | 5 | 3 |
|  |  |  |  |  |  |  |  |  | 14 | 19066305 | 24437922 | 2 | 1 |
|  |  |  |  |  |  |  |  |  | 14 | 24454989 | 25008758 | 4 | 2 |
|  |  |  |  |  |  |  |  |  | 14 | 25026780 | 34014124 | 2 | 1 |
|  |  |  |  |  |  |  |  |  | 14 | 34020584 | 35019099 | 3 | 1 |
|  |  |  |  |  |  |  |  |  | 14 | 35032112 | 99659371 | 2 | 1 |
|  |  |  |  |  |  |  |  |  | 14 | 99669020 | 104554044 | 3 | 2 |
|  |  |  |  |  |  |  |  |  | 14 | 104559552 | 106404410 | 5 | 3 |
|  |  |  |  |  |  |  |  |  | 14 | 106412532 | 107289541 | 2 | 1 |
|  |  |  |  |  |  |  |  |  | 15 | 20060562 | 40329520 | 1 | 0 |
|  |  |  |  |  |  |  |  |  | 15 | 40330693 | 42083843 | 1 | 1 |
|  |  |  |  |  |  |  |  |  | 15 | 42104339 | 42226255 | 3 | 2 |
|  |  |  |  |  |  |  |  |  | 15 | 42233908 | 42431236 | 2 | 1 |
|  |  |  |  |  |  |  |  |  | 15 | 42433382 | 42448658 | 3 | 2 |
|  |  |  |  |  |  |  |  |  | 15 | 42453954 | 90170232 | 1 | 1 |
|  |  |  |  |  |  |  |  |  | 15 | 90172832 | 90352611 | 3 | 2 |
|  |  |  |  |  |  |  |  |  | 15 | 90369148 | 95706135 | 1 | 1 |
|  |  |  |  |  |  |  |  |  | 15 | 95712020 | 97706696 | 2 | 1 |
|  |  |  |  |  |  |  |  |  | 15 | 97712183 | 97875116 | 1 | 0 |
|  |  |  |  |  |  |  |  |  | 15 | 97888564 | 101162066 | 1 | 1 |
|  |  |  |  |  |  |  |  |  | 15 | 101172258 | 102520957 | 3 | 1 |
|  |  |  |  |  |  |  |  |  | 16 | 97583 | 3170188 | 3 | 2 |
|  |  |  |  |  |  |  |  |  | 16 | 3173575 | 4356193 | 1 | 1 |
|  |  |  |  |  |  |  |  |  | 16 | 4367492 | 5223036 | 2 | 2 |
|  |  |  |  |  |  |  |  |  | 16 | 5247799 | 11203558 | 1 | 0 |
|  |  |  |  |  |  |  |  |  | 16 | 11225515 | 11607076 | 2 | 2 |
|  |  |  |  |  |  |  |  |  | 16 | 11618256 | 27751757 | 1 | 1 |
|  |  |  |  |  |  |  |  |  | 16 | 27761631 | 29814234 | 2 | 1 |
|  |  |  |  |  |  |  |  |  | 16 | 29818850 | 30134251 | 5 | 3 |
|  |  |  |  |  |  |  |  |  | 16 | 30147265 | 35213861 | 2 | 1 |
|  |  |  |  |  |  |  |  |  | 16 | 46438402 | 56899547 | 1 | 1 |
|  |  |  |  |  |  |  |  |  | 16 | 56901288 | 58079157 | 2 | 1 |
|  |  |  |  |  |  |  |  |  | 16 | 58092093 | 66240435 | 1 | 0 |
|  |  |  |  |  |  |  |  |  | 16 | 66246688 | 70624483 | 1 | 1 |
|  |  |  |  |  |  |  |  |  | 16 | 70630077 | 71174927 | 2 | 1 |
|  |  |  |  |  |  |  |  |  | 16 | 71188623 | 83949853 | 1 | 1 |
|  |  |  |  |  |  |  |  |  | 16 | 83953124 | 84211465 | 2 | 1 |
|  |  |  |  |  |  |  |  |  | 16 | 84212571 | 84270741 | 3 | 3 |
|  |  |  |  |  |  |  |  |  | 16 | 84296118 | 87349529 | 1 | 1 |
|  |  |  |  |  |  |  |  |  | 16 | 87350773 | 90292760 | 2 | 2 |
|  |  |  |  |  |  |  |  |  | 17 | 301 | 9846633 | 3 | 0 |
|  |  |  |  |  |  |  |  |  | 17 | 9856734 | 16321029 | 1 | 0 |
|  |  |  |  |  |  |  |  |  | 17 | 16322676 | 18286762 | 3 | 0 |
|  |  |  |  |  |  |  |  |  | 17 | 18314913 | 21195039 | 2 | 0 |
|  |  |  |  |  |  |  |  |  | 17 | 21198799 | 21321955 | 5 | 1 |
|  |  |  |  |  |  |  |  |  | 17 | 21356435 | 33502973 | 2 | 0 |
|  |  |  |  |  |  |  |  |  | 17 | 33504465 | 36614524 | 0 | 0 |
|  |  |  |  |  |  |  |  |  | 17 | 36617210 | 37558900 | 2 | 0 |
|  |  |  |  |  |  |  |  |  | 17 | 37561613 | 45745468 | 1 | 0 |
|  |  |  |  |  |  |  |  |  | 17 | 45750462 | 48834023 | 4 | 2 |
|  |  |  |  |  |  |  |  |  | 17 | 48843437 | 69919968 | 2 | 1 |
|  |  |  |  |  |  |  |  |  | 17 | 69928346 | 71189182 | 6 | 2 |
|  |  |  |  |  |  |  |  |  | 17 | 71192873 | 72710796 | 3 | 2 |
|  |  |  |  |  |  |  |  |  | 17 | 72733082 | 78846909 | 4 | 2 |
|  |  |  |  |  |  |  |  |  | 17 | 78854377 | 81194987 | 6 | 3 |
|  |  |  |  |  |  |  |  |  | 18 | 10813 | 52294 | 4 | 0 |
|  |  |  |  |  |  |  |  |  | 18 | 52880 | 76951258 | 1 | 0 |
|  |  |  |  |  |  |  |  |  | 18 | 76961923 | 78017073 | 2 | 0 |
|  |  |  |  |  |  |  |  |  | 19 | 266023 | 4844149 | 2 | 0 |
|  |  |  |  |  |  |  |  |  | 19 | 4847874 | 5307420 | 24 | 5 |
|  |  |  |  |  |  |  |  |  | 19 | 5320087 | 7431287 | 8 | 2 |
|  |  |  |  |  |  |  |  |  | 19 | 7437583 | 8675108 | 28 | 5 |
|  |  |  |  |  |  |  |  |  | 19 | 8741614 | 10057878 | 6 | 1 |
|  |  |  |  |  |  |  |  |  | 19 | 10070954 | 10265312 | 3 | 2 |
|  |  |  |  |  |  |  |  |  | 19 | 10267266 | 14200930 | 8 | 2 |
|  |  |  |  |  |  |  |  |  | 19 | 14231330 | 14412411 | 3 | 1 |
|  |  |  |  |  |  |  |  |  | 19 | 14425502 | 17396549 | 3 | 2 |
|  |  |  |  |  |  |  |  |  | 19 | 17397479 | 17655505 | 11 | 4 |
|  |  |  |  |  |  |  |  |  | 19 | 17660300 | 18119418 | 4 | 2 |
|  |  |  |  |  |  |  |  |  | 19 | 18119744 | 18174729 | 5 | 3 |
|  |  |  |  |  |  |  |  |  | 19 | 18180447 | 18950636 | 9 | 3 |
|  |  |  |  |  |  |  |  |  | 19 | 18956698 | 20913512 | 3 | 2 |
|  |  |  |  |  |  |  |  |  | 19 | 20918419 | 22559123 | 6 | 2 |
|  |  |  |  |  |  |  |  |  | 19 | 22585517 | 33044716 | 1 | 0 |
|  |  |  |  |  |  |  |  |  | 19 | 33054934 | 36038390 | 1 | 1 |
|  |  |  |  |  |  |  |  |  | 19 | 36041030 | 36259467 | 2 | 2 |
|  |  |  |  |  |  |  |  |  | 19 | 36266776 | 36643483 | 4 | 2 |
|  |  |  |  |  |  |  |  |  | 19 | 36667932 | 37397596 | 1 | 1 |
|  |  |  |  |  |  |  |  |  | 19 | 37399404 | 38109777 | 1 | 0 |
|  |  |  |  |  |  |  |  |  | 19 | 38127250 | 41497280 | 2 | 1 |
|  |  |  |  |  |  |  |  |  | 19 | 41510102 | 43237759 | 6 | 2 |
|  |  |  |  |  |  |  |  |  | 19 | 43243218 | 45253101 | 5 | 2 |
|  |  |  |  |  |  |  |  |  | 19 | 45287674 | 45911320 | 11 | 7 |
|  |  |  |  |  |  |  |  |  | 19 | 45912000 | 48692874 | 5 | 2 |
|  |  |  |  |  |  |  |  |  | 19 | 48697961 | 48785642 | 3 | 1 |
|  |  |  |  |  |  |  |  |  | 19 | 48800299 | 49132634 | 12 | 5 |
|  |  |  |  |  |  |  |  |  | 19 | 49133761 | 49141532 | 7 | 3 |
|  |  |  |  |  |  |  |  |  | 19 | 49143025 | 49206603 | 8 | 4 |
|  |  |  |  |  |  |  |  |  | 19 | 49214748 | 51455556 | 6 | 3 |
|  |  |  |  |  |  |  |  |  | 19 | 51463845 | 54314585 | 7 | 2 |
|  |  |  |  |  |  |  |  |  | 19 | 54327568 | 55879872 | 4 | 2 |
|  |  |  |  |  |  |  |  |  | 19 | 55895579 | 56215171 | 26 | 7 |
|  |  |  |  |  |  |  |  |  | 19 | 56220504 | 56717353 | 6 | 2 |
|  |  |  |  |  |  |  |  |  | 19 | 56717353 | 58858224 | 2 | 1 |
|  |  |  |  |  |  |  |  |  | 19 | 58858676 | 59118886 | 6 | 2 |
|  |  |  |  |  |  |  |  |  | 20 | 76962 | 2728558 | 4 | 3 |
|  |  |  |  |  |  |  |  |  | 20 | 2730400 | 3236015 | 7 | 6 |
|  |  |  |  |  |  |  |  |  | 20 | 3245006 | 3629686 | 3 | 2 |
|  |  |  |  |  |  |  |  |  | 20 | 3640467 | 3684022 | 9 | 8 |
|  |  |  |  |  |  |  |  |  | 20 | 3686436 | 3911270 | 6 | 5 |
|  |  |  |  |  |  |  |  |  | 20 | 3917072 | 24923660 | 3 | 2 |
|  |  |  |  |  |  |  |  |  | 20 | 24929947 | 25865400 | 4 | 4 |
|  |  |  |  |  |  |  |  |  | 20 | 25873122 | 30870339 | 5 | 3 |
|  |  |  |  |  |  |  |  |  | 20 | 30887184 | 33551218 | 4 | 3 |
|  |  |  |  |  |  |  |  |  | 20 | 33565480 | 33592474 | 9 | 9 |
|  |  |  |  |  |  |  |  |  | 20 | 33593674 | 43933163 | 3 | 2 |
|  |  |  |  |  |  |  |  |  | 20 | 43942676 | 44691350 | 5 | 4 |
|  |  |  |  |  |  |  |  |  | 20 | 44697431 | 47606043 | 3 | 3 |
|  |  |  |  |  |  |  |  |  | 20 | 47626847 | 47919672 | 5 | 3 |
|  |  |  |  |  |  |  |  |  | 20 | 47935629 | 50177662 | 3 | 3 |
|  |  |  |  |  |  |  |  |  | 20 | 50182797 | 53268372 | 7 | 4 |
|  |  |  |  |  |  |  |  |  | 20 | 53276418 | 55955567 | 5 | 3 |
|  |  |  |  |  |  |  |  |  | 20 | 55969697 | 56412867 | 7 | 5 |
|  |  |  |  |  |  |  |  |  | 20 | 56420820 | 60180124 | 3 | 3 |
|  |  |  |  |  |  |  |  |  | 20 | 60183964 | 62961724 | 7 | 6 |
|  |  |  |  |  |  |  |  |  | 21 | 9527088 | 14703726 | 4 | 2 |
|  |  |  |  |  |  |  |  |  | 21 | 14716293 | 33057344 | 2 | 2 |
|  |  |  |  |  |  |  |  |  | 21 | 33069118 | 35531416 | 4 | 3 |
|  |  |  |  |  |  |  |  |  | 21 | 35543613 | 39899525 | 3 | 2 |
|  |  |  |  |  |  |  |  |  | 21 | 39912560 | 42938226 | 4 | 3 |
|  |  |  |  |  |  |  |  |  | 21 | 42942620 | 45096206 | 5 | 4 |
|  |  |  |  |  |  |  |  |  | 21 | 45107518 | 47639492 | 7 | 6 |
|  |  |  |  |  |  |  |  |  | 21 | 47641700 | 48118698 | 4 | 4 |
|  |  |  |  |  |  |  |  |  | 22 | 16870802 | 18859378 | 4 | 0 |
|  |  |  |  |  |  |  |  |  | 22 | 18894046 | 20311131 | 7 | 0 |
|  |  |  |  |  |  |  |  |  | 22 | 20353879 | 22049783 | 4 | 0 |
|  |  |  |  |  |  |  |  |  | 22 | 22053880 | 28586674 | 2 | 0 |
|  |  |  |  |  |  |  |  |  | 22 | 28604583 | 31535995 | 4 | 0 |
|  |  |  |  |  |  |  |  |  | 22 | 31552702 | 37260123 | 2 | 0 |
|  |  |  |  |  |  |  |  |  | 22 | 37268257 | 38485064 | 4 | 0 |
|  |  |  |  |  |  |  |  |  | 22 | 38486738 | 50058338 | 2 | 0 |
|  |  |  |  |  |  |  |  |  | 22 | 50085205 | 51243277 | 1 | 0 |

Supplemental Table 5 Tumor purity and ploidy values calculated by tumor cell lines and clinical samples with different tumor purity

| ID | purity.pdt | purity.cal | ploidy.cal |
| --- | --- | --- | --- |
| HCC1143 | 0.80 | 0.83 | 3.47 |
|  | 0.60 | 0.59 | 3.47 |
|  | 0.50 | 0.50 | 3.44 |
|  | 0.40 | 0.39 | 3.43 |
|  | 0.30 | 0.29 | 3.35 |
|  | 0.25 | 0.24 | 3.27 |
|  | 0.20 | 0.20 | 3.31 |
|  | 0.10 | 1.00 | 2.12 |
| HCC1428 | 0.80 | 0.80 | 3.61 |
|  | 0.60 | 0.59 | 3.60 |
|  | 0.50 | 0.48 | 3.59 |
|  | 0.40 | 0.39 | 3.58 |
|  | 0.30 | 0.29 | 3.58 |
|  | 0.25 | 0.24 | 3.59 |
|  | 0.20 | 0.19 | 3.44 |
|  | 0.10 | 1.00 | 2.16 |
| HCC38 | 0.80 | 0.69 | 3.25 |
|  | 0.60 | 0.57 | 3.35 |
|  | 0.50 | 0.45 | 3.27 |
|  | 0.40 | 0.35 | 3.29 |
|  | 0.30 | 0.26 | 3.29 |
|  | 0.25 | 0.22 | 3.36 |
|  | 0.20 | 0.16 | 3.18 |
|  | 0.10 | 1.00 | 2.00 |
| clinical_sample1 | 0.47 | 0.55 | 2.60 |
|  | 0.31 | 0.36 | 2.60 |
|  | 0.16 | 0.19 | 2.30 |
| clinical_sample2 | 0.44 | 0.46 | 2.90 |
|  | 0.29 | 0.32 | 2.80 |
|  | 0.15 | 0.20 | 2.40 |
| clinical_sample3 | 0.27 | 0.32 | 2.00 |
|  | 0.18 | 0.20 | 2.00 |
|  | 0.09 | 0.09 | 2.10 |
| clinical_sample4 | 0.25 | 0.29 | 3.80 |
|  | 0.17 | 0.20 | 3.90 |
|  | 0.08 | 0.10 | 4.00 |
| clinical_sample5 | 0.33 | 0.34 | 1.90 |
|  | 0.22 | 0.23 | 1.80 |
|  | 0.11 | 0.14 | 2.00 |

Supplemental Table 6 Comparison of the tumor purity and ploidy predicted by GSA, PureCN, and ASCAT.

| ID | purity.pdt | GSA | | PureCN | | ASCAT | |
| --- | --- | --- | --- | --- | --- | --- | --- |
|  |  | purity.cal | ploidy.cal | purity.cal | ploidy.cal | purity.cal | ploidy.cal |
| HCC1143 | 0.8 | 0.83 | 3.47 | 0.84 | 3.62 | 0.8 | 3.91 |
|  | 0.6 | 0.59 | 3.47 | 0.62 | 3.66 | 0.65 | 3.85 |
|  | 0.5 | 0.5 | 3.44 | 0.5 | 3.66 | 0.61 | 3.17 |
|  | 0.4 | 0.39 | 3.43 | 0.45 | 3.59 | 0.53 | 3.03 |
|  | 0.3 | 0.29 | 3.35 | 0.32 | 3.71 | NA | NA |
|  | 0.25 | 0.24 | 3.27 | 0.32 | 2.2 | NA | NA |
|  | 0.2 | 0.2 | 3.31 | 0.32 | 2.25 | NA | NA |
|  | 0.1 | NA | NA | 0.2 | 2.31 | NA | NA |
| HCC1428 | 0.8 | 0.8 | 3.61 | 0.88 | 3.63 | 0.98 | 3.79 |
|  | 0.6 | 0.59 | 3.6 | 0.6 | 3.63 | 0.72 | 3.88 |
|  | 0.5 | 0.48 | 3.59 | 0.49 | 3.63 | 0.64 | 3.66 |
|  | 0.4 | 0.39 | 3.58 | 0.4 | 3.66 | NA | NA |
|  | 0.3 | 0.29 | 3.58 | 0.37 | 1.9 | NA | NA |
|  | 0.25 | 0.24 | 3.59 | 0.27 | 2.05 | NA | NA |
|  | 0.2 | 0.19 | 3.44 | 0.21 | 2.1 | NA | NA |
|  | 0.1 | NA | NA | 0.15 | 1.78 | NA | NA |
| HCC38 | 0.8 | 0.69 | 3.25 | 0.71 | 3.39 | 0.83 | 2.52 |
|  | 0.6 | 0.57 | 3.35 | 0.56 | 3.4 | 0.69 | 3.05 |
|  | 0.5 | 0.45 | 3.27 | 0.44 | 2.48 | 0.61 | 2.74 |
|  | 0.4 | 0.35 | 3.29 | 0.41 | 3.44 | 0.47 | 3.16 |
|  | 0.3 | 0.26 | 3.29 | 0.32 | 2.59 | NA | NA |
|  | 0.25 | 0.22 | 3.36 | 0.32 | 2.29 | NA | NA |
|  | 0.2 | 0.16 | 3.18 | 0.15 | 1.9 | NA | NA |
|  | 0.1 | NA | NA | 0.15 | 2.06 | NA | NA |
| clinical_sample1 | 0.74 | 0.74 | 3.00 | 0.80 | 3.18 | 0.87 | 3.65 |
|  | 0.44 | 0.46 | 2.90 | 0.50 | 3.18 | 0.58 | 3.14 |
|  | 0.29 | 0.32 | 2.80 | 0.33 | 3.21 | 0.45 | 2.35 |
|  | 0.15 | 0.20 | 2.40 | 0.17 | 2.31 | NA | 2.08 |
| clinical_sample2 | 0.45 | 0.45 | 2.00 | 0.46 | 2.17 | 0.65 | 1.84 |
|  | 0.27 | 0.32 | 2.00 | 0.32 | 2.15 | 0.48 | 1.67 |
|  | 0.18 | 0.20 | 2.00 | 0.22 | 2.17 | NA | 2.21 |
|  | 0.09 | NA | NA | 0.15 | 2.12 | NA | 4.19 |
| clinical_sample3 | 0.42 | 0.42 | 3.90 | 0.52 | 4.26 | NA | NA |
|  | 0.25 | 0.29 | 3.80 | 0.39 | 4.19 | NA | NA |
|  | 0.17 | 0.20 | 3.90 | 0.34 | 3.88 | NA | NA |
|  | 0.08 | NA | NA | 0.22 | 3.81 | NA | NA |
| clinical_sample4 | 0.54 | 0.54 | 1.90 | 0.45 | 2.04 | 0.64 | 1.44 |
|  | 0.33 | 0.34 | 1.90 | 0.31 | 2.02 | 0.97 | 3.19 |
|  | 0.22 | 0.23 | 1.80 | 0.22 | 2.05 | NA | 2.37 |
|  | 0.11 | NA | NA | 0.15 | 2.03 | NA | 4.26 |
